# Individualized temporal patterns dominate cortical upstate and sleep depth in driving human sleep spindle timing

**DOI:** 10.1101/2024.02.22.581592

**Authors:** Shuqiang Chen, Mingjian He, Ritchie E. Brown, Uri T. Eden, Michael J. Prerau

**Author notes:** Corresponding author: Michael J. Prerau.

## Abstract

Sleep spindles are critical for memory consolidation and strongly linked to neurological disease and aging. Despite their significance, the relative influences of factors like sleep depth, cortical up/down states, and spindle temporal patterns on individual spindle production remain poorly understood. Moreover, spindle temporal patterns are typically ignored in favor of an average spindle rate. Here, we analyze spindle dynamics in 1008 participants from the Multi-Ethnic Study of Atherosclerosis using a point process framework. Results reveal fingerprint-like temporal patterns, characterized by a refractory period followed by a period of increased spindle activity, which are highly individualized yet consistent night-to-night. We observe increased timing variability with age and distinct gender/age differences. Strikingly, and in contrast to the prevailing notion, individualized spindle patterns are the dominant determinant of spindle timing, accounting for over 70% of the statistical deviance explained by all of the factors we assessed, surpassing the contribution of slow oscillation (SO) phase (∼14%) and sleep depth (∼16%). Furthermore, we show spindle/SO coupling dynamics with sleep depth are preserved across age, with a global negative shift towards the SO rising slope. These findings offer novel mechanistic insights into spindle dynamics with direct experimental implications and applications to individualized electroencephalography biomarker identification.

## 1. INTRODUCTION

Sleep spindles, a subset^1,2^ of transient neural oscillatory bursts (typically lasting 0.5 to 1.5 seconds) during non-rapid eye movement (NREM) sleep, are visually observed as ∼ 10 – 16 Hz fluctuating waveforms in the sleep electroencephalogram (EEG) in humans^3–6^. Since their initial discovery in 1935^4^, sleep spindles have been of great interest due to their association with memory consolidation^7–9^, learning^10–12^, neurological disorders (such as Alzheimer’s disease^13–15^, schizophrenia^16–19^, epilepsy^20,21^, and Parkinson’s disease^22,23^), as well as noted changes during natural aging^24–27^.

Numerous studies have demonstrated that spindle activity dynamically and continuously evolves over time and is mediated by a variety of intrinsic and extrinsic factors including sleep stage^5,28^, slow oscillation activity (0.5 – 1.5 Hz)^25,29–34^, hippocampal ripples^29,33,35,36^, and infra-slow oscillations (< 0.1 Hz)^5,37–40^. The timing and rate of occurrence of spindles is also influenced by homeostatic and circadian factors, chronotype, biological sex, and IQ, as well as the biophysical properties of thalamic reticular nucleus and thalamocortical neurons, their synaptic connectivity and their modulation by ascending arousal systems^5^.

Despite these known dynamics, the relative influences on the moment-to-moment likelihood of a spindle event occurring at a specific time are not well-characterized. Moreover, standard analyses almost universally report average spindle rate (known as spindle density) over fixed stages or time periods—thus ignoring timing patterns completely. Recent studies, however, have started to assess temporal patterns of spindles. Antony et al. reported the refractoriness of spindles by visualizing the distribution of inter-spindle lags and highlighted the importance of spindle timing in optimal memory reactivation^40^; Boutin and Doyon proposed the idea of temporal clusters of spindles by defining the “train of spindles” as a group of at least 2 consecutive spindles interspaced by less than 6 seconds^39^, and based on this criterion, Champetier et al. further studied how spindle temporal clustering changes over age and their relevance for memory consolidation^38^. Those attempts to understand spindle temporal patterns highlight the importance of spindle dynamics in various functional roles. While promising, these results require further verification, as substantial bias may be imposed through the ad hoc thresholding, smoothing, and other assumptions used in these approaches^1^. Furthermore, it is yet to be seen whether these putative spindle clusters may arise from ultradian dynamics, such as periodic variability of sleep depth^41,42^, or are emergent properties of other intrinsic factors. Thus, it is vital to continue to explore these questions using a rigorous statistical framework that can explicitly model temporal dynamics.

There currently exist many open questions related to spindle temporal dynamics. Are spindle events independent of each other? If not, how do past spindle events influence the next spindle? How consistent is the spindle timing pattern within participant night-to-night? Is spindle/SO coupling stationary over the night or does it evolve over depth-of-sleep? What are the contributions of these factors to spindle generation, and is there any interaction between them? What heterogeneity in these individualized features is observed in populations as a function of demographics and age? Answering these types of questions is crucial to gaining insight into the underlying mechanisms of spindle activity.

The goal of this paper is to address these fundamental questions using objective quantitative analysis of spindle data. To do so, we develop a formal statistical framework to characterize the temporal patterns of spindles and disentangle the moment-by-moment influences of multiple factors on the instantaneous spindle rate. This approach is based on the theory of point processes. Point processes are mathematical models that describe discrete events that occur in space or time such as the arrival of customers to a store, the location of trees in a forest, or earthquake occurrences. Over the past several decades, point processes have been successfully applied in various fields including seismology^43^, neuroscience^44^, finance^45^, and recently in the characterization of sleep apnea event timing^46^. To our knowledge, however, point processes have not been used to explicitly model sleep spindle timing patterns. One well-known, but limited, class of point processes is the Poisson process, which is characterized by a single, constant rate of occurrences and assumes independence between event times (i.e., no patterns). Mathematically, computing the spindle density is equivalent to fitting a Poisson process model to the data. A general point process allows for non-stationary rates and dependency between events.

In this study, we characterize the instantaneous dynamics of spindles and quantify the influence of factors of interest across a large, multi-ethnic, cross-sectional dataset of 1008 human participants using a point process framework. We first evaluate how past spindle event history modulates the current spindle rate. This allows us to create a quantitative description of each patient’s individualized spindle patterns, as well as to make population-level comparisons across gender and age. We then investigate SO/spindle coupling as a dynamic process, quantifying how coupling changes with sleep depth and how it evolves over aging. Finally, we assess whether history dependence is related to SO phase.

## 2. RESULTS

### 2.1. Study overview: Modeling mechanisms of temporal influences with point processes

Spindle activity is a dynamic process that is governed by multiple temporal processes. Characterizing whether and how various intrinsic and extrinsic factors modulate spindle dynamics can provide insight into the mechanisms underlying spindle timing and form a rigorous basis for assessing demographic differences. Our study targets the following fundamental questions: (1) How do past spindle events impact the timing of future events? (2) How do spindles dynamically couple with slow oscillations over sleep depth? (3) What is the interplay of spindle history and SO in explaining spindle dynamics? Answering these questions provides insight into the mechanisms underlying spindle generation, the different functional roles of multiple factors in modulating spindle dynamics, and the variability in spindle patterns within individuals and across demographic populations.

Standard quantitative approaches to sleep spindle analysis have limitations that make answering these questions difficult. Sleep spindle analyses traditionally report the average spindle rate over a given sleep stage. As the configuration of events within the period does not affect the metric, subjects with drastically different temporal patterns are indistinguishable under this approach given similar average rates. Spindle metrics computed over stages are also sensitive to the demonstrably high inter-scorer variability in human manual scoring^47^ or variability across the multitudinous array of automated methods^1,48^.

Moreover, typical spindle/SO coupling analyses independently detect slow waves and spindles and throw out any events that do not mutually coincide. This imposes a bias on the estimated influence of SO on spindle production, as all uncoupled events are ignored. Thus, the degree to which SO phase influences spindle timing may potentially be overstated.

While it may be possible to directly explore these questions experimentally at the circuitry level in animal models, given the accessibility of large cross-sectional repositories of human sleep EEG data^49^, we can also address these questions through a statistical modeling approach. Here, we develop a point process – generalized linear model (GLM) framework to explicitly model spindle temporal patterns and quantify the influence of multiple factors (e.g., sleep stage, slow oscillation activity, etc.) on spindle activity^44,50,51^. A general point process can be defined in terms of a *conditional intensity function*, *λ*(*t*|*H*_*t*_), which describes an “instantaneous rate” at time *t*, given *H*_*t*_, the history of past events up to, but not including time *t*. The conditional intensity can then be expressed as a function of the covariates that influence spindle events. To model spindle dynamics, we express the conditional intensity of spindle events as a function of sleep stage, slow oscillation power (SOP, a surrogate of continuous sleep depth), slow oscillation phase, past spindle events, as well as the interactions between these factors. In doing so, we develop a formal statistical framework to determine quantitatively which factors affect the moment-by-moment spindle rate, providing a basis for the identification and analysis of spindle biomarkers. See Methods 4.2 - 4.4 for the overview of point process modeling and the specific details of our spindle GLMs.

We fit these models to a dataset with two-night sleep EEG recordings that included 17 participants, as well as a large cross-sectional dataset (the Multi-Ethnic Study of Atherosclerosis (MESA) dataset) with single-night recordings for 1008 participants. For each polysomnography record, we identified time-frequency peaks^1^ in the sigma range (12 to 16 Hz), aligning with the frequencies of traditional definitions of “fast” spindles and fit the models with their associated covariates (see Methods). Model performance was assessed, and hypothesis tests were conducted to determine the influence of each factor. We compared results at the level of individual participants (within/across nights) to the level of the entire MESA population, as well as across different gender and age groups.

### 2.2. Spindle timing patterns: past spindle activity influences the next spindle

Temporal patterns of events are very common in many physical processes and can reflect important aspects of function and features of underlying mechanisms. For example, consider firing patterns in neural spiking activity. Neurons typically enter an absolute refractory period after discharging an action potential, during which neurons are not excitable, which may also be followed by periods of increased excitation^52^. Given the relationship between the refractory period and excitatory periods, numerous neural firing patterns can be generated, such as bursting, that may additionally vary based on their intrinsic properties and the way they respond to different stimuli^53,54^. These varied firing patterns can play different critical roles in neural information coding, processing, and synchronization^55,56^. Using our framework, we can formally address the degree to which spindle timing exhibits stereotyped patterns. To do so, we incorporate *history dependence* into our point process models, which can be defined as the influence of past spindle events on the current spindle. To do so, we create a model where the likelihood of a spindle at each time depends on the time since the previous event. We visualize this effect using a *history modulation plot*, which answers the question: How much more likely is an event now given that the most recent event occurred x seconds ago?

Figure 1 shows a history modulation plot for an individual from the MESA dataset. The history curve (solid blue) shows the multiplicative effect on the current spindle rate for each time lag, with a 95% confidence interval (dashed). Portions of the modulation curve below 1 indicate a decreased rate, or refractory period during which few or no events occur, and portions above 1 indicate a period of increased rate of events. To explicitly quantify features of individualized timing properties in a more interpretable manner, we define features of the history modulation curve (see Methods 4.8 for details).

**Figure 1:**
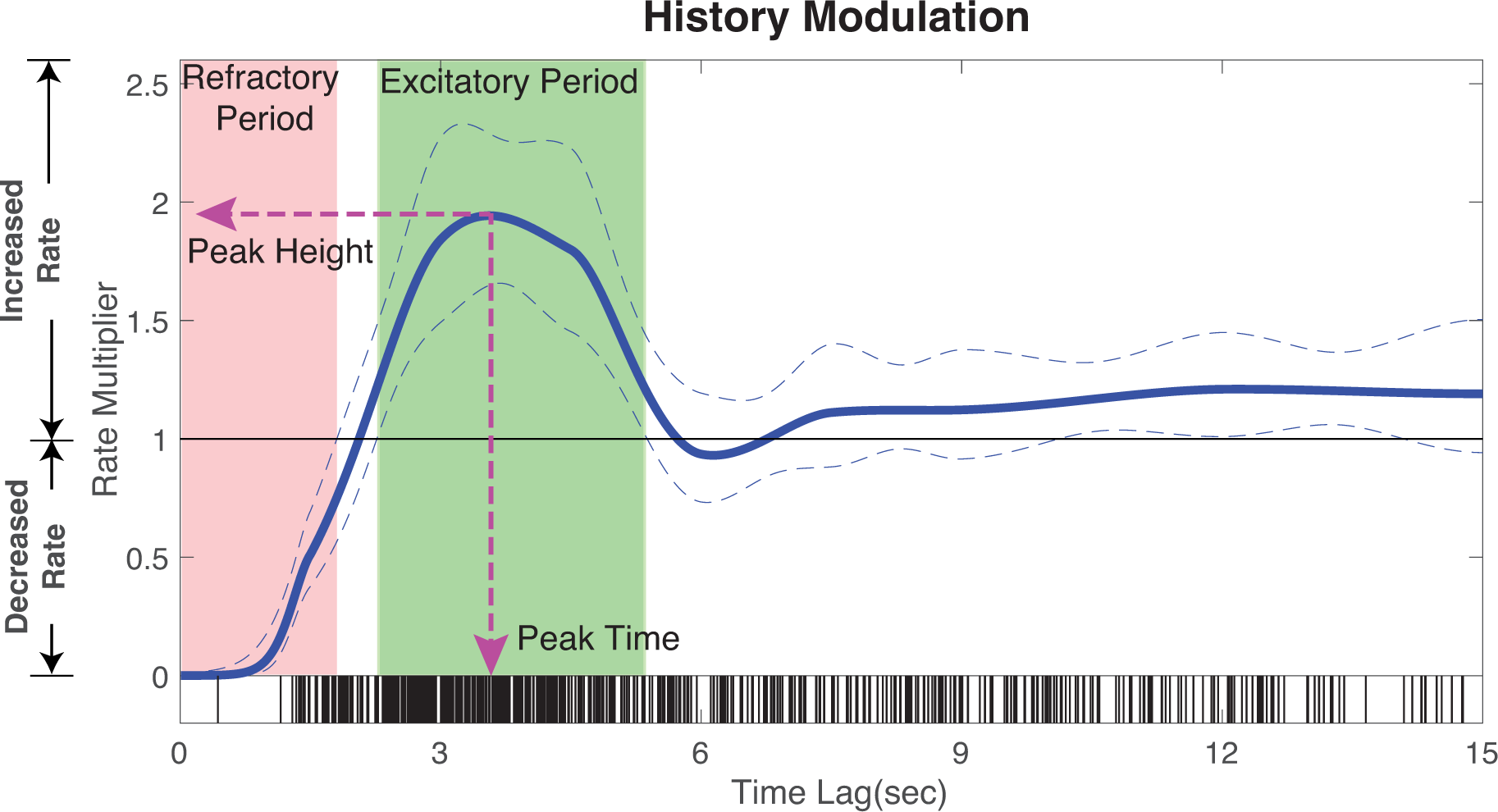
History modulation curve captures spindle timing patterns. Using a participant from the MESA dataset, the history modulation plot (solid blue curve) shows the multiplicative effect on event rate based on the time since the last event. 95% confidence intervals are shown as dashed lines. We summarize the history curve by defining the refractory period, excitatory period, peak height, and peak time.

Specifically, we quantify the following features:

- Refractory period: The duration after a spindle during which activity is significantly suppressed. It is quantified as the portion of the history modulation curve significantly less than 1.
- Excitatory period: The duration after the refractory period during which activity is significantly enhanced. It is quantified as the portion of the history modulation curve significantly greater than 1.
- Excitatory peak height: The degree to which the excitation is maximally enhanced.

It is quantified as the maximum of the rate multiplier in the excitatory period.

- Excitatory peak time: The lag at which the maximal excitation occurs. This suggests the most probable interval between spindles. It is quantified as the lag corresponding to the excitatory peak height.

Together, these features provide an enhanced level of interpretability to the history modulation curves. For this participant, the curve indicates a significant reduction in rate between 0 s and ∼1.8 s after the previous event (refractory period, red region), followed by a period between lags of ∼2.3 s and ∼5.3 s where the activity is significantly enhanced (excitatory period, green region). The curve reaches the maximum at around 3.5 s (peak time) with a modulation value close to 2 (peak height), indicating that at this lag, the spindle rate is nearly double the baseline rate of the current stage. Then the modulation curve fades back to 1 at a ∼6s lag, which suggests that events that have occurred more than 6s in the past will no longer have any significant effect on the current spindle rate.

### 2.3. Participants with similar spindle density can exhibit different timing patterns

As stated previously, a substantial shortcoming of spindle density is the inability to account for differences in temporal patterns. Consequently, drawing inferences based solely on spindle density may provide an incomplete or inaccurate description of the underlying dynamics. We illustrate this issue in Figure 2, where we show six participants from the MESA dataset that have nearly identical spindle density (N2 rate: 5 ± 0.5 events/min), yet exhibit temporal patterns with marked structural differences.

**Figure 2:**
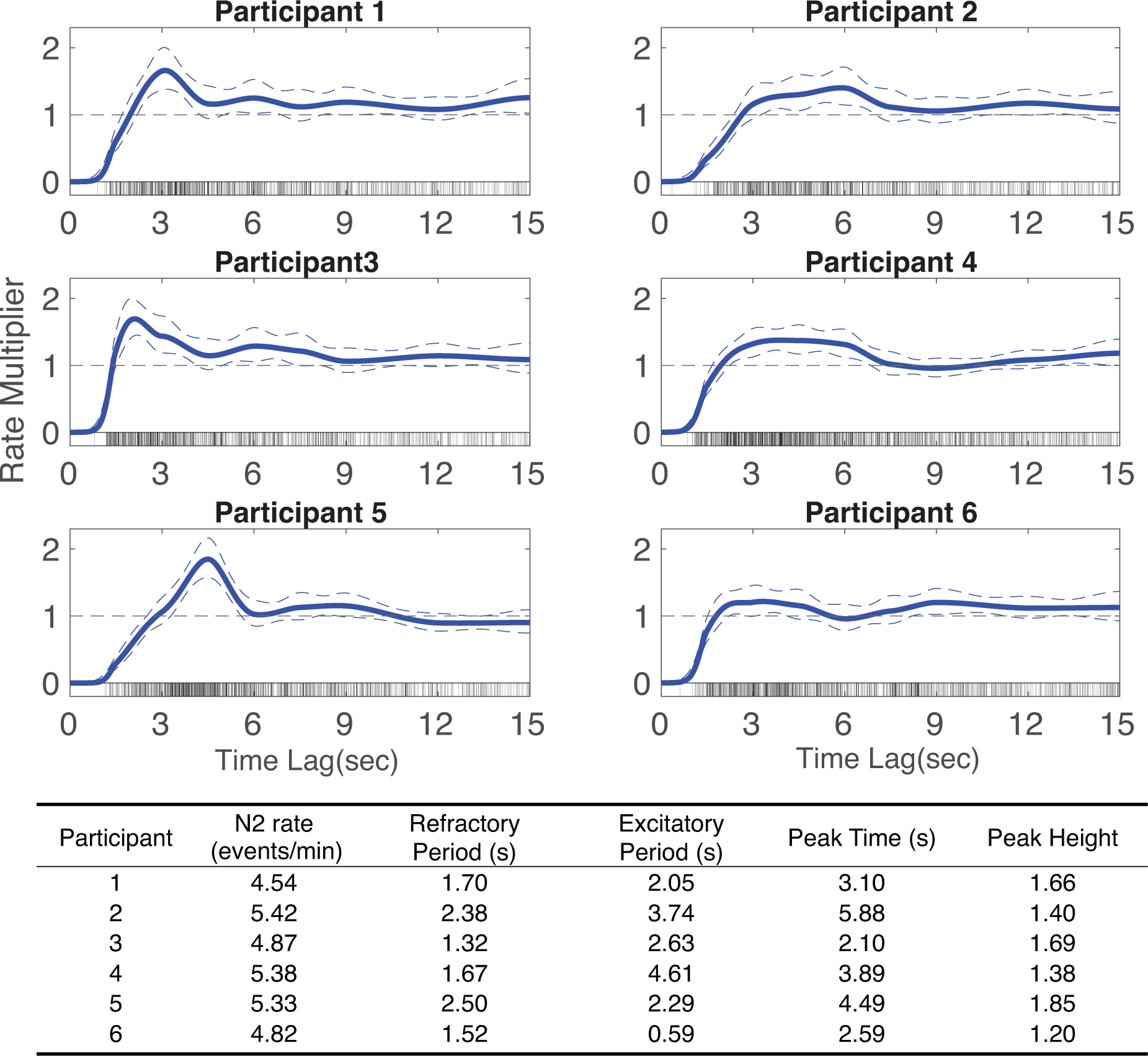
Participants with similar spindle densities can have drastically different history dependence structures. Six participants are shown with very similar spindle rates (N2 rate: 5 ± 0.5 events/min) who possess different history dependence architectures. History dependence curves are shown as solid blue lines and 95% confidence intervals are shown as dashed blue lines. In the bottom panel, a summary table summarizes show history features of each participant, including refractory period, excitatory period, peak time, and peak height.

All participants share the same general form of history modulation: a refractory period followed by a peak of increased propensity of events, which then decreases to 1. However, the structure of these features varies from subject to subject. For example, the tall, narrow peaks exhibited by participants 1,3 and 5 suggest more periodic trains of events. However, they exhibit vastly different refractory periods and peak times, suggesting different levels of periodicity. Whereas the broad, low peaks exhibited by participants 2 and 4 suggest larger variability and randomness in event patterns. Additionally, in examining contrasts between participant 4 and 6, we see participant 6 has a shorter excitatory period and peak height, suggesting even less predictable events than participant 4. Overall, variation in history dependence structure reflects remarkably heterogeneous temporal patterns, despite similar densities.

### 2.4. Spindle timing patterns are consistent night-to-night but heterogeneous in populations

Characterizing spindle history dependence structure helps us assess the heterogeneity across participants and consistency within participants across multiple nights. Figure 3a shows four healthy control participants from a small dataset (N = 17) from Wamsley et al.^19^ with recordings from two consecutive nights that have similar N2 spindle density (∼11 events/min). Again, we see substantially different levels of variability and periodicity of spindle timing patterns across subjects, yet night-to-night variability is small within subjects.

**Figure 3:**
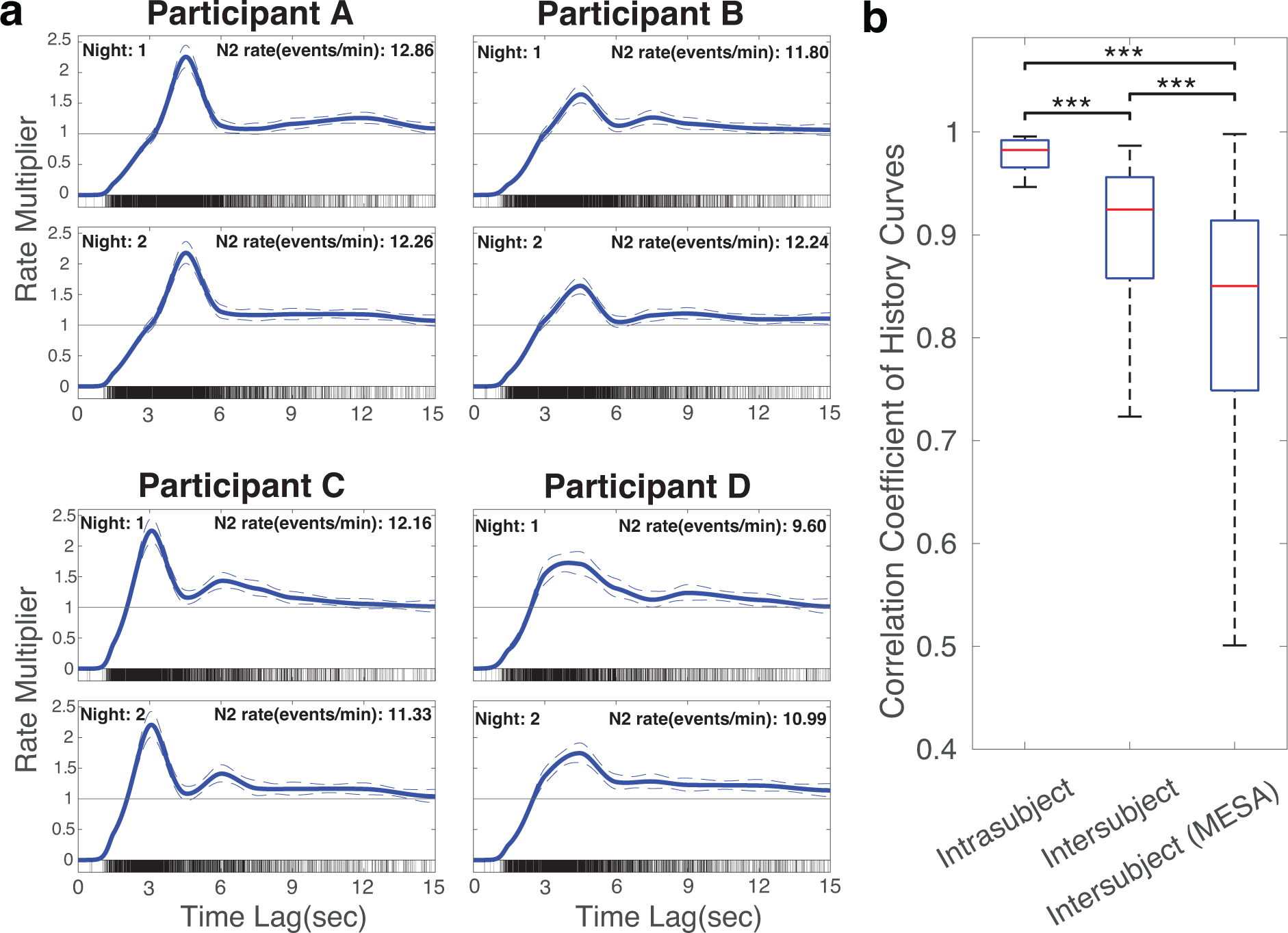
History dependence creates unique “temporal fingerprints” for each participant’s spindle activity, which suggest a basis for characterizing spindle phenotypes. (a): History curves of four participants with 2-night sleep. (b): Boxplots of the correlation coefficients between history curves for the same participant across nights (intrasubject) and between participants (intersubject) in Wamsley dataset (17 participants from 2 nights), as well as between participants in the MESA dataset (1008 participants). While the history dependent structure is drastically different between participants, night-to-night variability within participants is small.

To quantify the consistency within subjects and heterogeneity in populations, we report the intersubject and intrasubject correlation coefficient of history modulation curves in the small dataset (Wamsley) contrasted with the intersubject variability across the large cross-sectional MESA dataset (Figure 3b). In the smaller cohort with two available nights, we observe the distribution of intrasubject (same participant across two nights) and intersubject (different participants, first night) correlations. The mean intrasubject correlation coefficient of 0.98, with a significantly higher than mean intersubject variability of 0.90, with a much smaller variability in the intrasubject distribution. The heterogeneous MESA population (1008 participants) shows the lowest correlation coefficient (0.81) and largest variability. Notably, correlation coefficients in all groups are highly positive in general (mean values > 0.8), indicating that despite vast heterogeneity among populations, most participants share a consistent pattern that commonly involves a refractory period followed by an excitatory period. Overall, history dependence creates an individualized temporal signature of spindle activity and demonstrates strong within-subject stability across nights. This implies history dependence structure can potentially serve as a quantitative basis for future spindle phenotype exploration and clinical biomarker development.

### 2.5. Spindle temporal patterns show robust gender and age differences in a large heterogeneous population

Given the ability to quantify the individualized spindle timing patterns, we can characterize the variability in history features over populations. We summarize the results for the MESA cohort in Figure 4.

**Figure 4:**
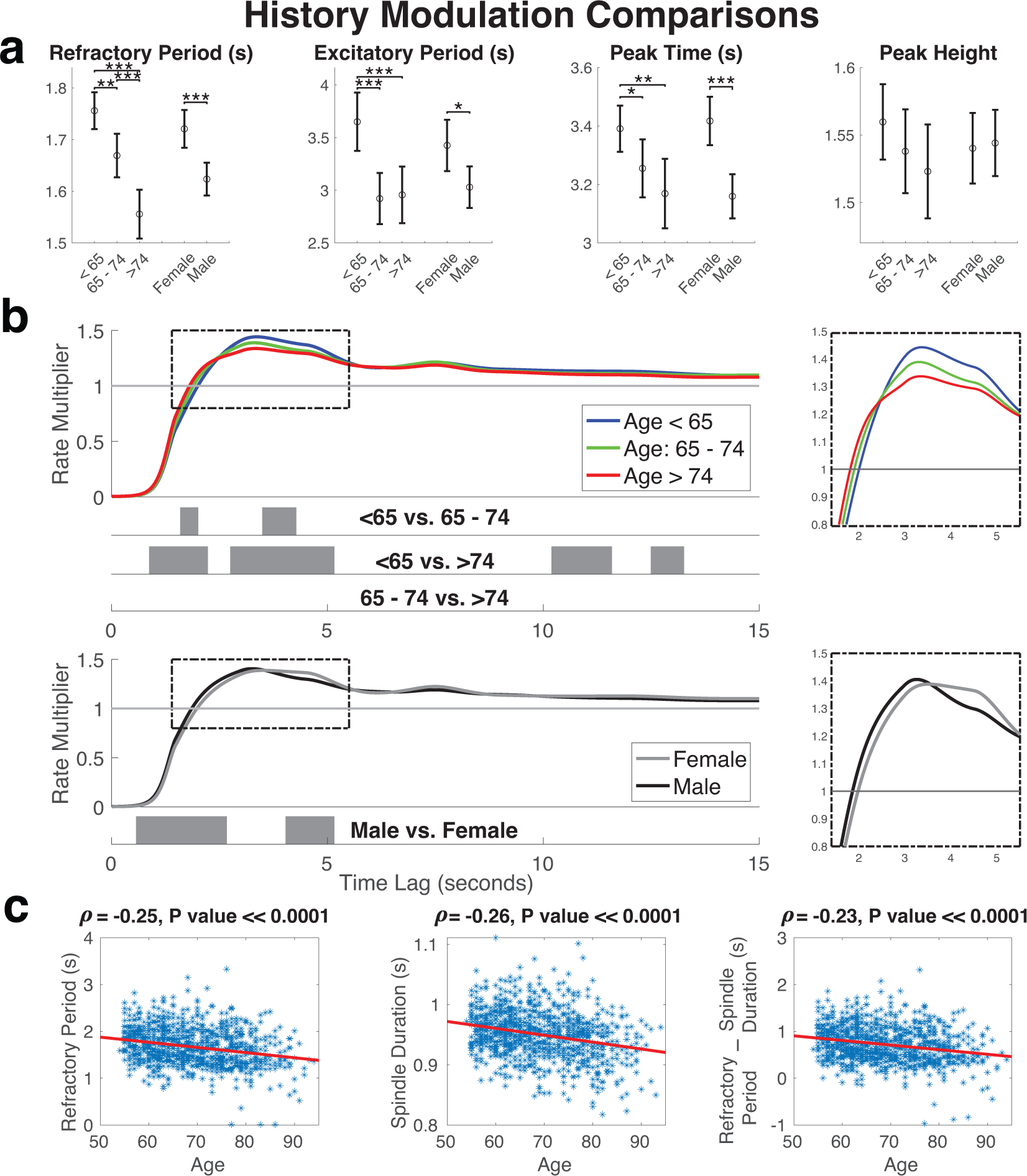
History dependence features show robust demographic differences. **(a)**: Comparisons of all summary statistics of history modulation across groups. (*,**,*** denote p values < 0.05, < 0.01, < 0.001, respectively for t-tests comparing group means, not corrected for multiplicity). (b): The mean history modulation curves with significant regions (gray) using global permutation tests (Significance level: .05), The dashed box on the right shows a magnification highlighting key features. (c): Scatter plots of refractory period, spindle duration, and refractory period - spindle duration vs. age suggest a reduction in the refractory period with age that cannot be explained by shorter duration spindle events. The fitted linear regression line is shown in red, and correlation correfficients and their P values are reported for each plot.

Figure 4a compares features of history curves among different demographic groups, and Figure 4b shows the average history modulation curves as a function of gender and age groups, with corresponding regions where these curves differ significantly (gray regions). The refractory period becomes significantly shorter with age. One possibility is that this reduction in the refractory period is due to the shorter spindle duration in older people as reported in multiple previous studies^6,26,57–59^. However, after subtracting the spindle duration, we find that the refractory period continues to exhibit a persistent and statistically significant negative correlation with aging (Figure 4c). This suggests that independent of duration, there are significant changes in the intrinsic timing properties of spindles, contributing to reduced refractory period with age.

We also find a significant reduction in the duration of the excitatory period along with a lower peak height (though not statistically significant) with aging. This suggests that spindle patterns become less periodic and more variable with age, which provides additional evidence of age-related impairment of sleep spindle dynamics^25,38^.

Distinct differences in history curves are also observed among gender groups, where males exhibit significantly shorter refractory periods, excitatory periods, and earlier peak times. Moreover, spindle patterns in aging females generally have similar properties to younger males, which is a common observation in sleep data^26,60^. The correlation between these extracted history properties and other key spindle morphology features is shown in Supplementary Figure 1.

### 2.6. SO/spindle coupling shows a continuous negative phase shift with increasing sleep depth

While characterizing spindle history dependence allows us to better characterize temporal patterns, many other factors also impact spindle occurrence. Perhaps the most notable feature thought to influence spindle timing is the slow oscillation (SO) phase, which is thought to reflect cortical up/down states. It has been widely reported that fast spindles tend to occur at a preferential phase at the peak of SO (upstate)^25,31,33,61–63^. It is the prevailing practice to detect spindles and slow waves separately and then select only those that cooccur temporally, rejecting all other detected events. Consequently, the average SO/spindle coupling phase is typically computed during the N3 stage, where co-occurrence is more likely^25,31^. Alternatively, both N2 and N3 stages may be analyzed, but spindles that occur within a specific timeframe of slow waves will be considered^33,36,62^. These approaches raise several concerns: First, coupling analysis is most often focused on N3 given the definitional increased prevalence of SOs, yet targets fast spindles, which are most predominant in N2^5^. Furthermore, by discarding all events that do not cooccur, analyses make the implicit assumption that SOs are 100% necessary for spindle production, while also introducing selection bias via both spindle and SO detection^1^. Finally, most coupling analyses are broken up into discretized sleep stages to account for stage-dependent coupling, which assumes stationarity within stages and restricts the analysis of coupling dynamics across the continuum of sleep depth.

We address sleep stage-specific phase coupling by building a model of spindle activity as a function of sleep stage, SO phase, and phase-stage interaction terms. This model allows us to capture stage-specific preferred phases. This approach also provides us with an estimate of statistical uncertainty about the preferred phase and allows us to perform hypothesis tests to determine whether SO/spindle coupling depends on sleep stage. In Figure 5b, the solid red lines show the preferred phase in each sleep stage and the dashed red lines show the 95% confidence interval for a single participant. We observe that the preferred phase shifts from the peak of the SO (∼ 0 phase) to the rising phase of SO (∼ -*π*/4 phase) as this participant moves from stage N2 to N3. Sleep stage has a statistically significant influence on SO/spindle coupling (*χ*^2^ test, p ≪ .0001) for this participant. Note that since there are very few events detected in N1 stage (Figure 5a), this creates a vast uncertainty in phase preference (confidence interval spans -*π* to *π*), suggesting that N1 does not have a statistically relevant preferred phase.

**Figure 5:**
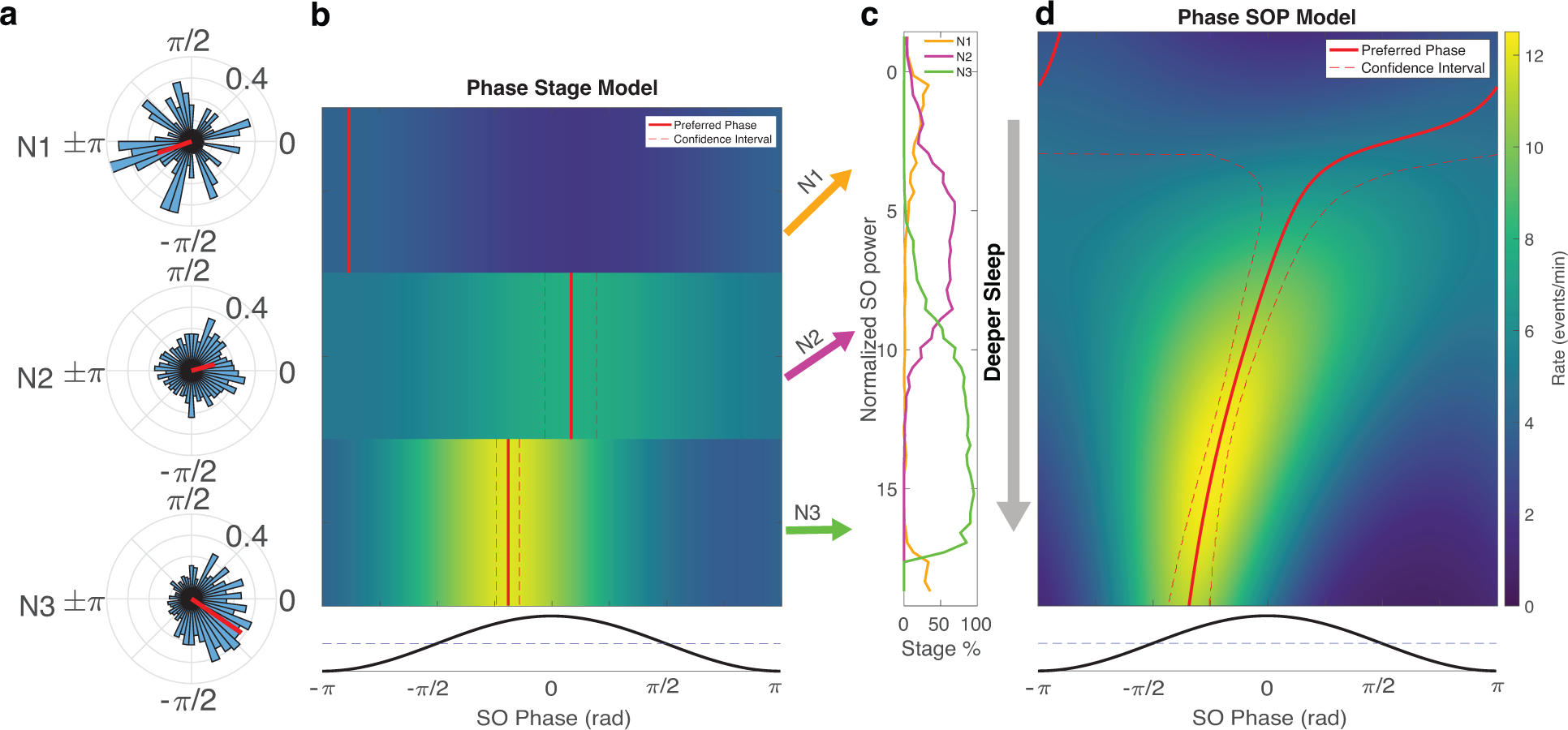
Analysis of spindle/SO coupling as a function of sleep depth for a single participant. (a): Polar histograms of spindle phase in stages N1, N2 and N3, with mean phase (red arrow); (b): Colormap of the fitted spindle rate as a function of SO phase (x-axis) and sleep stage (y-axis). The red solid and dashed lines indicate the estimated preferred phase and its confidence interval from the point process model; (c): The percentage of time the participant was in scored sleep stages N1, N2, and N3 as a function of the normalized SO power; (d): Colormap of the fitted spindle rate as a function of SO phase (x-axis) and SO power (y-axis). The red solid and dashed lines indicate the estimated preferred phase and its confidence interval; These results reveal that the spindle/SO coupling shifts to a more negative phase as this participant moves to the deeper sleep.

As sleep is a continuous, dynamic process, the natural extension is to examine how SO/spindle coupling is influenced by sleep depth when considered as a continuum. Slow oscillation power (SOP) has been shown to be an objective and robust metric of continuous sleep depth^2,64^. Thus, we replace the discrete stages in the previous model with SOP. For the same participant, we see the preferred phase shows a continuous gradient from the SO peak to the SO rising phase as the participant falls deeper into sleep (Figure 5d). This aligns with the result from the discrete stage model but provides a more precise characterization of the within- and across-stage coupling dynamics, revealing the continuous evolution of SO/spindle coupling with depth of sleep.

### 2.7. SO/spindle coupling shows a negative phase shift across all sleep depths in aging

We examined the spindle phase dependency across the MESA cohort. *χ*^2^ tests show 839 out of 1008 participants (83.23%) had significant dependence between spindles and SO phase. We further assessed sleep depth dependent coupling across these 839 participants and compared the results over three age groups in MESA cohort (age range: 54 - 94), as well as in a much younger Wamsley group (age range: 26 - 45). Figure 6 a and b show the distribution of the preferred phase (gray polar histograms) and the mean preferred phase (red line) across each sleep stage and SOP category. As people move to deeper sleep, we see a consistent and statistically significant shift (Watson-Williams tests, p ≪ .0001) of the preferential phase from the peak of the SO to the rising slope of the SO.

**Figure 6:**
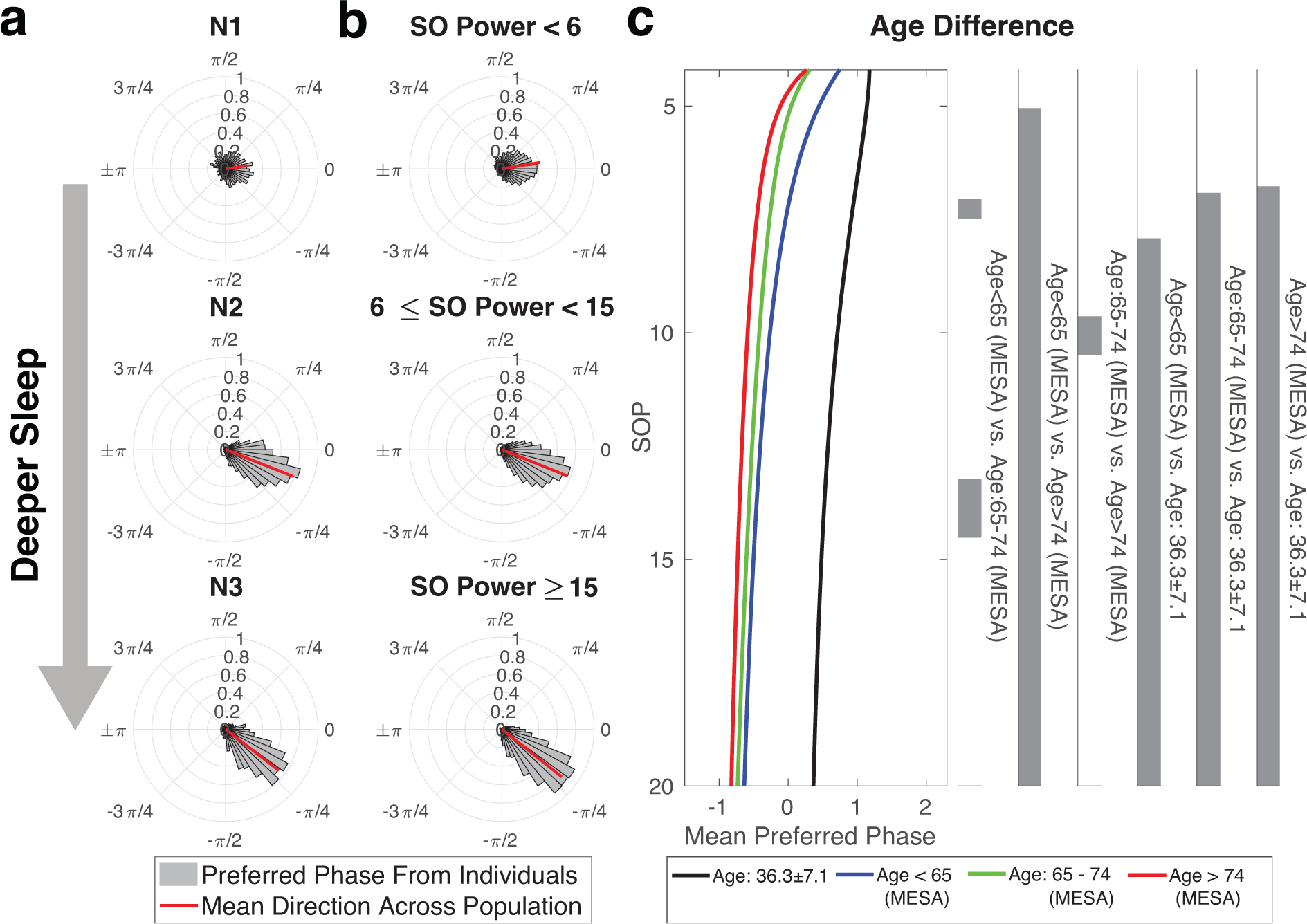
Population trends reveal shifts in preferential phase over sleep depth. Sleep spindles tend to occur at the up state of SO (peak of slow oscillation, ∼ 0 phase of cosine curve). As participants in this population move to the deeper sleep, their preferential phase shifts towards the down state (trough) of the SO. The results are shown for both the discrete (**a**) and continuous (**b**) sleep depth metrics. Gray polar histograms show the distributions of preferred phase across the population, with the red line showing the mean preferred phase. (**c**): Mean phase shift curves from the phase-SOP model are shown across three age groups in the MESA dataset (blue: < 65, green: 65 – 74, red: > 74), as well as in the Wamsley dataset (black, mean age: ∼ 36) with significance reginons determined by global permutation tests for all combinations (Significance level: .05).

Though evidence of stage-dependent phase coupling has previously been reported in studies from Cox et al^28^, due to the small sample size (11 participants were included in N3, and 14 participants in N2) and their method for computing preferred phase, the authors did not have sufficient statistical power to identify a significant difference in fast spindle/SO coupling between N2 and N3. Our results show the same direction of phase shift from N2 to N3, but also provide a statistically powerful way to detect small shift patterns that would not be identifiable through simple summary statistics. We also fit a *phase-SOP model* to characterize the SO/spindle coupling as a function of continuous sleep depth.

Figure 6c shows the mean phase coupling curve across three age groups in the MESA dataset and the younger group from Wamsley et al.^19^ We observe a global phase shift across all depths of sleep as a function of age, reflecting the negative shift previously reported in the literature^25^. This is critical because it demonstrates that this shift is not solely due to the age-dependent differences in sleep depth. Rather, we see similar trajectories with increasing age across all sleep depth levels.

### 2.8. Among multiple factors, history is the most important component for predicting spindle timing

We have demonstrated that spindle rate can be modulated by the history of past spindle events and coordinated with slow oscillation activity over sleep depth. We now ask: How do these individual factors interact in shaping spindle dynamics, and how much does each contribute to spindle timing? To address these questions, we construct a series of point process models that contain combinations of these factors. Specifically, we fit single-factor models of spindle conditional intensity using history, phase, stage (discrete sleep depth), and SOP (continuous sleep depth) alone, as well as multi-factor models that included history, phase, and sleep depth (one with scored stage & stage-phase interaction, and one with SOP & SOP-phase interaction). Using the population data, we then assessed the degree to which each factor contributes and the degree to which the factors provide independent information about spindle timing.

Figure 7 compares fits for all proposed models for a single participant from MESA. Figure 7a shows detected spindle events (red dots) across the full night on a multitaper spectrogram of the EEG along with the corresponding hypnogram. Below, are the estimated intensities for the single-factor (stage, SOP, and phase) and multi-factor (stage/SOP + phase + history) models for a single NREM bout of ∼1h duration, which is further expanded into a 1.5-minute time segment from N2 sleep (Figure 7b).

**Figure 7:**
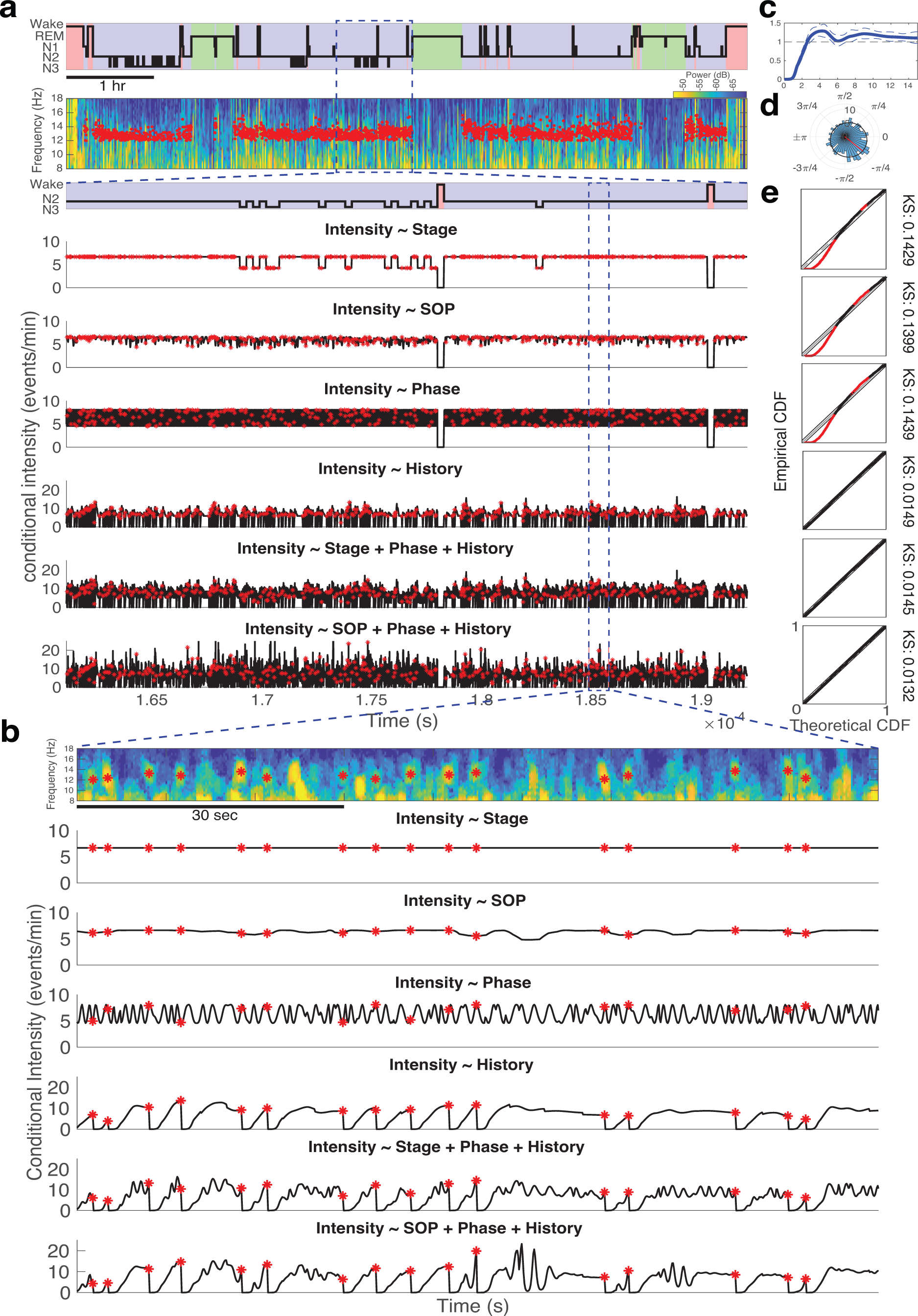
Model estimates from sleep spindle data for a single participant and night. **(a)**: From top to bottom, hypnogram for the entire night; EEG multitaper spectrogram with red dots indicating detected spindles; Sleep stage time series; Model fits for multiple models over a period of about 1 hour. **(b)**: Magnification of spectrogram and model fits from panel (a) over 1.5-min. (c): History modulation curve for this participant for the model using only history as a factor; (d): polar histogram of spindle phase overlaid with the estimated conditional intensity as a function of phase from the model with only SO phase as a factor (black); the preferred phase estimate is shown as a red arrow; (e): KS plots and statistics for each model; regions exceeding the 95% test bounds are marked in red.

We first examine the single-factor models. In the *stage model*, the spindle rate is a discrete, stepwise function that only depends on the current stage. The event rate is highest in N2 (∼ 6.7 events/min) and goes down in N3 (∼ 4.3 events/min). As expected, there is a low rate in N1 (∼ 3.5 events/min) and a rate of zero in REM and wake periods (in which spindles were not detected). Using the continuous sleep depth model of the SOP, we observe a similar structure to the stage model, but with more variability within-stage due to continuous shifts in SOP. The single-factor phase model fluctuates between 4.6 and 8 events/min as a function of phase. The spindle intensity (black) as a function of phase is overlaid on the polar histogram of the spindle SO phase in Figure 7b, as a reference. From the polar histogram and the 1.5-min segment, it is clear that while generally spindles tend to occur in the up-state of SO, which is consistent with many previous studies^25,32,36^, many spindles also occur at other phases and even phases corresponding to the down-state.

We next examine the single-factor history model. The fitted modulation curve (Figure 7c) shows a typical structure: a refractory period followed by a peak at ∼ 4 seconds and gradual return towards 1. We can see the structure of this history dependence reflected in the temporal structure of the conditional intensity in Figure 7b. Immediately after each spindle event the intensity drops to zero and remains there through the refractory period. The intensity rises during the excitatory period, and often a new spindle is generated as the intensity approaches its peak. This can lead to a sequence of high density spindles. If no spindle occurs during the excitatory period, the intensity gradually declines to its baseline level where it settles until the next event. As multiple events arrive sequentially, the history of multiple past events can interact, with excitatory and inhibitory periods combining constructively or destructively depending on spindle timing.

It is important to note that for the single-factor sleep depth and phase models (with the exception of Wake, REM, and N1 in which spindles are not typically found) the range of the conditional intensity is fairly narrow from a minimum of about 4.5 events/min to a maximum of 8 events/min. The lack of any periods when the intensity approaches zero suggests that phase and sleep depth indicate when spindles are more probable as a general rule but are not particularly informative about individual spindle timing. In particular, for the phase model this suggests that spindles can still occur 180° from the preferred phase of the SO. In contrast, the range of the spindle intensity for the history model is around 3 times greater, ranging from a minimum near-zero to a maximum around 25 events/min. This suggests that the times of past events predict spindle times and non-spindle times with a much higher certainty than the other factors.

This improvement is quantified in Figure 7e, which shows Kolmogorov-Smirnov (KS) plots used to evaluate the model goodness-of-fit^44,65^. The KS plot describes the degree to which the observed inter-spindle times are predicted by the model. A well-fit model will have a KS plot that follows the 45-degree line and stays within the global test bounds (gray region). Any excursion outside of these test bounds indicates a significant lack of fit (marked by red). The KS statistic is also reported for each model, which measures the largest deviation between the KS plot and the y = x line. For this participant, the stage, SOP, and phase models fail the KS test (KS statistics = 0.1429, 0.1399 and 0.1439, respectively). However, the history model, even without other predictors, leads to a KS plot that remains within the test bounds (KS statistic = 0.0149).

We also fit multi-factor models, to capture the combined influence and interaction between multiple factors on the occurrence of spindle events. In particular, the multiple-factor model that includes SOP as a continuous measure of sleep depth (KS statistic = 0.0132) outperforms the model that uses discrete sleep stage (KS statistic = 0.0145). Figure 7b shows that the SOP model and its interaction with phase recovers spindle dynamics related to the SO that are otherwise ignored by the discrete stage model.

### 2.9. History dependence is pervasive across participants and the dominant contributor to spindle dynamics

To assess the generalizability of these findings in the population, we fit each of these models and assessed the contributions of each factor across 1008 participants in the MESA population.

We compared the KS test pass rate for all models. The model that included only stage as a factor had a KS pass rate of 14.29%. This was less informative than the model with the SOP as a continuous measure of sleep depth, for which the KS pass rate was 19.21%. The model with only SO phase had a KS pass rate of 16.07%. The relatively low pass rate of these single factor models stand in contrast with the model that includes only history dependence, which had a KS pass rate of 72.02%. Combining these factors in the *SOP-Phase-History Interaction* model yields a KS test pass rate of 98.71%.

For this full model with multiple factors, a deviance analysis was performed to compare the contribution of each factor. Model deviance extends the concept of model variance in linear regression to point process data. By comparing nested models, we can compute the percentage of the deviance in the full model explained by adding each subsequent factor (see Methods for computation of deviance explained). We find that among all factors, the history component contributes the most (the mean fraction of the full deviance explained across the population was 72.11%, with a 95% confidence interval (CI) of [71.15%, 73.07%]), as compared to stage, SOP and phase components that have mean fractional deviance explained at 16.58% (CI [15.90%, 17.47%]), 18.33% (CI [17.23%,19.42%]), and 13.60% (CI [12.88%,14.31%]), respectively.

In addition, we evaluate the pervasiveness of spindle history dependence within the general population. After fitting history models to the 1008 participants from the MESA cohort, *χ*^2^tests reveal that 991 out of 1008 participants (98.31%) exhibit a statistically significant history effect using just one night of data, suggesting that history dependence is a near-universal feature of spindle activity.

### 2.10. History and slow oscillation phase have independent influences on spindle dynamics

Given the large proportion of information gained by the inclusion of spindle history dependence, one mechanistically critical question arises: Do history dependence and SO phase carry separate of redundant information about spindle timing? We address this question by computing the synergy index, a common measure in neural modeling derived from information theory^66–69^. A value of −1 indicates that two neurons provide completely overlapping information (totally redundant), 0 indicates entirely independent information, and 1 indicates that each neuron contributes little information on its own, but together they yield substantially more information (totally synergistic). The specific formula for this index is provided in Methods. Figure 8 shows the distribution of the synergy index across 1008 MESA participants. The distribution of the synergy index is tightly concentrated around 0, which suggests that history and phase provide mutually independent information in modulating spindle dynamics for every participant in the study.

**Figure 8:**
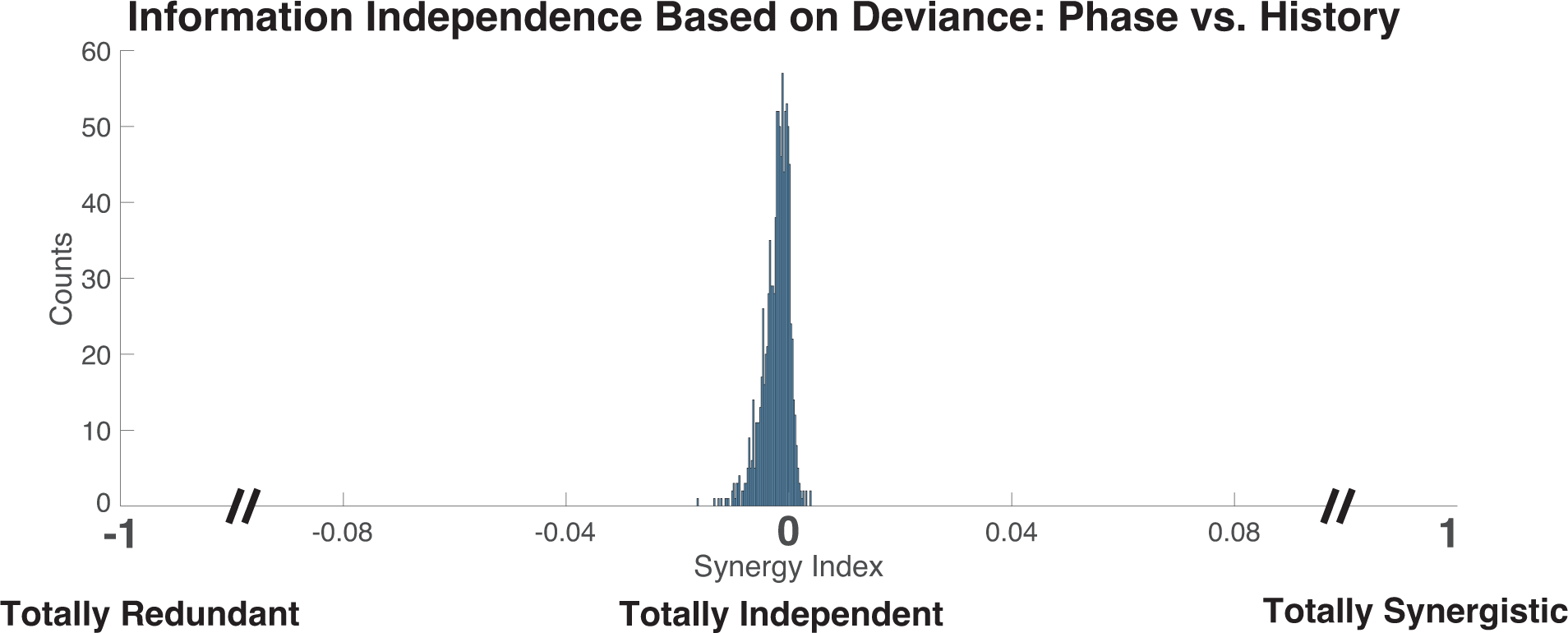
History dependence and slow oscillation phase provide independent information about spindle dynamics. The distribution of the synergy index across the MESA dataset is concentrated around 0, indicating that history and SO phase provide independent information.

As an alternative approach, a mean correlation matrix of fitted parameters across the MESA cohort from the multi-factor model with stage, phase, and history (without interactions) is shown in Supplementary Figure 2. We can see that SO phase and history components have almost no correlation with each other, which corroborates their independent roles in modulating spindle dynamics.

## 3. DISCUSSION

Sleep spindle activity is a dynamic process in which the rate of events is simultaneously governed by multiple factors and their interactions, the intricacies of which are not fully understood. In this work, we developed a quantitative approach to characterize the factors contributing to spindle timing, measuring the relative influence of different sources, as well as their interactions. We model the instantaneous rate of spindle events as a function of any combination of the factors influencing it, including sleep stage, past spindle history, slow oscillation activity, and their interactions. This approach provides a means of explicitly testing which factors influence spindle generation and timing within a rigorous statistical framework, augmenting the inferential power of existing experimental data. Our results suggest new and informative features of spindle patterning indicative of natural heterogeneity and potentially informative about pathophysiological processes. The comparative assessment of these models underscores the significance of individual factors contributing across multiple time scales, combining short-term influences of SO phase, slower dynamics related to sleep depth, and event history, which has long and short-term influences.

### 3.1. Spindle activity is history dependent

Our results suggest that spindle production is history dependent, such that the timing of a spindle is influenced by the timing of previous events. This effect is pervasive, with nearly all participants sharing a similar general history modulation structure: starting with a refractory period in which events do not occur, followed by an excitatory period, in which there is an increased likelihood of events (Figure 1). The structure of these history dependent factors creates patterns in spindle timing, variation in which creates differences between individuals. Like other features of spindle-like transient oscillations^2,28^, the history dependent structure exhibits strong night-to-night consistency for individuals, (Figure 3), with intersubject heterogeneity, even for participants with similar N2 spindle density (Figure 2, Figure 3).

We also observe robust differences in history dependent structure across gender and age groups (Figure 4). In particular, we show that females are more robust to alterations in history with the aging process, which aligns with previous findings showing men experience worse sleep disruption and NREM sleep impairment than women^26,60,70^. We also show that the spindle refractory period shortens with aging independently of age-related reductions in spindle duration. The reduction in the excitatory period and peak height with aging suggests a loss of spindle rhythmicity and an increase of randomness in the elderly. This corroborates with the previous study showing decreased spindle clustering with aging but provides a complete and unbiased characterization of spindle temporal structure without relying on hard thresholds to define the temporal clustering level^38^. These altered history features provide another perspective on age-impaired hippocampal-dependent memory and learning^16,26,27^. Overall, these findings show the potential for this model framework to address specific epidemiological questions and mechanistic hypotheses in future applications.

### 3.2. History dependence is the dominant driver of spindle timing, far above phase and sleep depth

The inclusion of history dependence in our models has the effect of drastically improving the predictability of subsequent events and model goodness-of-fit. In the MESA population, ∼98% exhibit a statistically significant history effect using just one night of data. The importance of history is evident in the example in Figure 7, for which the addition of the history component greatly increases confidence in spindle event timing. Substantial increases in predictability are common when history dependence is included in our point process models^44,46^.

History dependence is by far the factor that is most predictive of spindle timing when considering the relative influences of history, SO-phase, and sleep depth. Specifically, the deviance analysis on the full population shows that over ∼70% of the deviance explained by our full model is explained by the history dependence component, compared to only ∼14% for phase, and ∼16% for sleep depth. These results are surprising, given widespread reports suggesting cortical up/down states as a crucial factor in modulating spindle dynamics^5^. While history dependence, as modeled by this approach, reflects the influence of multiple neurophysiological mechanisms, we have explicitly shown that the information encompassed by history dependence is independent of the information provided by the SO phase.

### 3.3. The assumption of spindle/slow wave cooccurrence imposes bias on mechanistic interpretation

Our analysis and modeling approach provide a number of results that challenge the existing perspective on spindle production. From the polar histogram of individuals (Figure 4a, Figure 7d), we see while spindles preferentially occur in the up-state of SO, many spindles also appear at other phases or even in the SO down-state. Figure 7b suggests spindle production is a complex and multifaced process, involving multiple factors at different time scales, each of which contributes to providing enhancement or suppression of spindle likelihood at any given point in time. Thus, it is the combination of multiple factors that provide various windows of opportunity for spindle activity to occur. The dominance of history dependence from deviance analysis implies that intrinsic factors of spindle production such as refractory/excitatory periods may override extrinsic triggering factors such as SO-phase. These provide a dramatically different view than mechanistic models that suggest that up-states are explicitly necessary for the production of spindles.

Several possibilities could contribute to this alternative viewpoint on spindle production. In particular, the pervasive practice of discrete spindle and slow-wave cooccurrence in coupling analyses for both EEG and intracranial studies could deeply bias hypotheses. Methodologically, this approach consists of independently detecting spindles and slow waves, then restricting the analysis to only those detected spindle events that sufficiently temporally overlap with detected slow wave events. These waveform detection algorithms have been shown to be highly variable in their results^1,33,36,48^ with ad hoc parameter settings that can create rarity assumptions that underselect events with low amplitude but identical morphology^1^. For this reason, studies often only examine SO/spindle phase coupling in deep NREM sleep (N3 - N4), so that they may find enough slow waves that pass the detection threshold. Consequently, low cooccurrence rates have been explicitly noted during N2^31^. However, this is suboptimal since fast spindle activity is definitionally maximal in N2 sleep^5^, so the majority of spindles throughout the night are being ignored. Selection bias is further exacerbated by the common practice of further restricting the analysis to only the detected spindles of the largest amplitude (e.g., top 25%).

Beyond these methodological issues, forced cooccurrence raises theoretical questions. By removing any slow waves and spindles that do not cooccur, this approach imposes the assumption that spindles without slow waves do not exist. Given a scenario in which SO-phase intermittently or weakly contributes to spindle production, as suggested by the population model analysis results, cooccurrence would definitionally induce a stronger importance of phase coupling. Thus, the absolutism of spindle/slow-wave coupling requires further study and justification.

Crucially, the cooccurrence assumption directly impacts the ability to observe history dependence by drastically reducing the chance of observing runs of consecutive spindles over consecutive slow waves. Without these runs, it is impossible to see short-term history processes like refractory periods and excitatory periods, as cooccurring spindles will rarely be close enough temporally to interact with previous events. Consequently, thus far, the timing of spindles and their phase coupling have had to be examined separately, as cooccurrence eliminates any dependence on previous events or history. In using this new approach, we are now able to measure the competing influences of these processes on spindle timing.

Alternatively, given the myriad of complexities of electrophysiological volume conduction in signals, referencing and montage choices, and spatial specificity of neural activity, it is nearly certain that we are observing the superposition of signals from multiple sources in any EEG recording, and to some lesser extent in intracranial recordings. Thus, it is unclear whether analysis methods can adequately match the “correct” slow waves and spindles when we observe a down-state spindle or a slow wave devoid of spindle activity. However, given the potential to observe both spindle and slow wave events from other spatial regions at a given electrode, we would expect history dependence and phase coupling to be equally corrupted. Therefore, the incorrect matching of events or the presence of superfluous events alone cannot explain the dominance of history dependence nor its independent information contribution from phase coupling.

### 3.4. Dynamic spindle/SO coupling over sleep depth

Our study provides a conclusive demonstration of significant spindle/SO coupling shifts dynamically in a continuum with sleep depth (sleep stage & SOP) in a large dataset. This agrees with existing studies reporting phase coupling in N2 and N3^5,25,31,61,71^, as well as those suggesting differences but lacking the statistical power to show the difference^28^. Moreover, we expand upon previous studies^25,31,61^ showing a fixed phase shift towards the rising edge of the SO with age, by showing that phase coupling dynamics as a whole is largely preserved with age, and that the whole continuum is globally shifted earlier.

### 3.5. Implications for spindle generation mechanisms

Studies in experimental animals provide some clues about the physiological mechanisms underpinning our observations. Sleep spindles recorded from cortical sites are generated through the interplay of neuronal activity in GABAergic thalamic reticular nucleus (TRN) neurons, especially the major population containing the calcium binding protein parvalbumin^72–74^ and glutamatergic thalamocortical (TC) neurons which in addition to projecting to the cortex also send collateral projections that feedback and excite TRN neurons^5,75^. During wakefulness, both TRN and TC neurons are relatively depolarized due to the excitatory action of ascending neuromodulatory inputs. When these inputs are withdrawn during NREM sleep, the ensuing hyperpolarization brings TRN and TC neurons into the appropriate voltage range where low-threshold calcium channels (Cav3.3 and Cav3.2 in TRN neurons, Cav3.1. in TC neurons) can become active. Activation of low-threshold calcium channels leads to burst discharge of TRN neurons and a compound inhibitory postsynaptic potential (IPSP) in TC neurons which hyperpolarizes them and removes inactivation of low-threshold calcium channels. Once the IPSP decays, these channels are activated, leading to a burst discharge of TC neurons, resulting in activation of cortical neurons and activation of TRN neurons, restarting the cycle of the spindle oscillation. Various biophysical and synaptic connectivity and kinetic properties of TRN and TC neurons affect the waxing and waning of spindles, their amplitude, frequency and duration^5^.

In contrast to the mechanisms underlying the generation of spindles themselves, less research has focused on the mechanisms regulating temporal spindle patterns. Over long timescales (hours), spindle occurrence is regulated by homeostatic and circadian mechanisms^5,76^. Recent studies in mice have also described an infraslow (∼50 s) cycle regulating the likelihood of spindle occurrence which is controlled by the activity of locus coeruleus noradrenaline neurons^77,78^. Other neuromodulatory inputs from serotonergic raphe neurons and cholinergic basal forebrain and brainstem neurons may also play a role, related to their control of the depth of sleep and transitions between sleep stages^75^. However, the timescale of these slow and ultraslow modulations is longer than the 3-20 s period following a detected spindle we have focused on here. *In vitro* studies have investigated the possible ionic mechanisms underlying the refractory period in TRN and TC neurons following spindle-like burst activity. In TRN neurons, a study in ferret brain slices *in vitro* described a slow afterhyperpolarization of TRN neurons following repetitive bursting caused by sodium and calcium-dependent potassium currents^79^. In TC neurons, *in vitro* studies in rats and cats^80,81^ have described a pronounced afterdepolarization following repetitive burst discharge due to prolonged activation of hyperpolarization and cyclic nucleotide gated cation (HCN) channels which generate the so-called H-current. This afterdepolarization will prevent further burst discharge of TC neurons, which is required for spindle generation by increasing inactivation of low-threshold calcium channels and taking the membrane potential of TC neurons out of the range in which they can be activated. The afterdepolarization is caused by increased intracellular calcium due to entry through low-threshold (Cav3.1) calcium channels, leading to calcium-activated production of cyclic AMP, which activates HCN channels^82–85^. Large scale computational models of thalamocortical activity underlying sleep spindles also point to calcium modulation of HCN channels as an important controller of spindle refractoriness^86^. Following the refractory period, we describe a variable period of enhanced spindle likelihood. The mechanisms responsible for the rebound excitation are less clear but may involve changes in the inactivation of low-threshold calcium currents that occur during the refractory period, short-term synaptic plasticity of TRN-TC, TC-TRN or cortical inputs to TRN, as well as alterations in cortical input to TRN. Factors which affect the synchronization of TRN and TC discharge may also affect the amplitude and form of the rebound excitation. Large-scale computational models may be helpful in understanding the relative importance of these factors^87^.

The factors responsible for interindividual differences in the temporal history of spindles and the effect of biological sex, age, and disease remain to be uncovered, but as described in the previous paragraph are likely due to differences in the biophysical properties of TRN and TC neurons, their intracellular sodium and calcium dynamics, their synaptic connectivity, kinetics and plasticity, with each other and with the cortex^5,88^, as well as interactions with ascending neuromodulatory and GABAergic inputs to the TRN and TC neurons^75,89^. Our study, which provides a robust unbiased framework to describe the temporal history of spindles, sets the stage for such mechanistic/genetic studies in humans and potentially also in animals, which can uncover the basis for inter and intraindividual differences and the effect of learning on these dynamics.

#### Extensions and future directions

Spindle timing exhibits individualized patterns across the population yet shows strong stability across nights, suggesting a quantitative fingerprint that can potentially serve as a diagnostic and therapeutic target for improving sleep-dependent memory and learning (Figure 2, Figure 3). Future work to understand how spindle patterns relate to neurological disease processes would shed light on the mechanistic roles of spindles in neuropathology and provide new insights into treatment efficacy.

Evidence has shown that spindles tend to cluster at an infraslow timescale (∼ 50 seconds)^37–39^. In this work, we modeled short-term history dependence structure (within 15 seconds), but this model component can be extended to test and quantify long-term dependence structure. Our current model assumes that participants have a single history dependence profile that is consistent across the whole night of sleep. A natural extension is to include a stage-history interaction term or a SOP-history interaction term to capture changing patterns of spindles with sleep depth.

Researchers have also reported a three-way coupling between spindles, SOs, and hippocampal ripples^29,33,35,36^. However, current practice uses a triple detection, triple cooccurrence approach, which further compounds the issues previously discussed, and can likewise impose extreme bias on some critical questions, such as the causality in SO-triggered spindles. While we are not able to detect ripples from normal sleep EEG, this framework can be applied to intracranial data to understand the mechanisms of such higher order associations.

Researchers have also found that spindle features, including spindle duration, frequency, and amplitude, are highly correlated with demographic differences and neurological disorders^1,5,16–18,57,90,91^. We have previously developed a class of marked point process models for neural spiking data, which model the temporal patterns and factors influencing spikes with different waveform structures^92,93^. These marked point process models can be extended to address issues inherent in spindle detection^1,2,48^ and target specific scientific questions related to spindle morphology. Marked point process models will also enable us to target the relationship between different classes of spindle events^2^ without defining ad hoc frequency cutoffs.

To conclude, we show that spindle temporal pattern factors explain over ∼70% of the statistical variability, in comparison with only ∼14% explained by cortical up/down state. These fingerprint-like timing patterns exhibit remarkable individualization across participants yet demonstrate strong consistency within participants across nights, with distinct differences across age and gender groups. Mechanistically, our study provides novel insights into spindle production, indicating that sleep depth sets a baseline rate that is predominantly modulated by intrinsic temporal patterns characterized by the refractory period and excitatory periods, with lesser and independent mediation by SO phase. Methodologically, we offer a statistically-principled framework to tackle fundamental spindle questions in a rigorous way, which could facilitate studies of sleep-dependent memory consolidation, characterize spindle abnormalities in neuropsychiatric disorders, and contribute to the development of new sleep biomarkers.

## 4. METHODS

### 4.1. Data description

We analyzed data from two datasets. The first is the Multi-Ethnic Study of Atherosclerosis (MESA) which contains a large cross-sectional population of participants^49,94^. The polysomnography (PSG) data from MESA was obtained from the National Sleep Research Resource (www.sleepdata.org). EEG data from electrode C4-M1, with a sampling frequency of 256 Hz, were analyzed. The sleep stages were labeled as Wake, rapid eye movement sleep (REM), and non-REM stages 1–3 (N1, N2, and N3), and each participant was labeled as one of three age groups (age < 65, age: 65 – 74, age > 75). We limited our analysis to 1008 participants (male/female: 523/485, age: mean 68.74 ± 8.95) that had high EEG quality (EEG signal good >= 95% of sleep time) and a spindle rate of at least 1 event/min in N2 or N3 stage.

The other dataset was from a previously published study by Wamsley et al.^19^ that included two-night sleep EEG recordings. The control group had 17 healthy participants who were screened to ensure no previous record of mental illness, family history of schizophrenia spectrum disorder, or psychoactive medication use. EEG data from electrode C3 (linked mastoids), with a sampling frequency of 100 Hz, was analyzed.

### 4.2. Developing a point process framework for spindle events

To establish a point process model, we first define the counting process for spindle events, denoted as *N*(*t*), which counts the total number of spindles that occur up to and including time *t*, where *t* ∈ (0, *TST*] and *TST* corresponds to the total sleep time (excluding all wake periods during sleep). The spindle density is defined as *N*(*TST*)/*TST*.

The conditional intensity function^50^ is a measure of the instantaneous probability of a spindle event at each time,

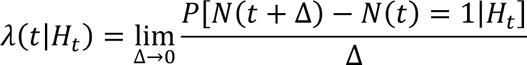

where *λ*(*t*|*H*_t_) is the conditional intensity at time t, which depends on *H*_t_, the past history of events up to, but not including time *t*.

For a specific set of times *s*_1_ < *s*_2_ < ⋯ < *s*_n_ over the interval [0, *T*], the likelihood of observing spindles at those times is^50^:

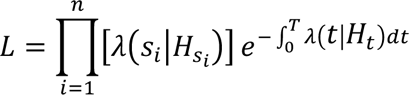

Therefore, the conditional intensity defines the probability distribution of observing a spindle event at each instant, which can be used to compute the likelihood of any pattern of events over the course of the night.

### 4.3. Model identification

Defining a point process model of spindle events involves expressing the conditional intensity, *λ*(*t*|*H*_t_), as a function of the variables that influence these events. One well-studied class of point process models fits into the broader statistical framework of generalized linear models (GLMs)^44,50,51^. Point process GLMs typically express the log of conditional intensity as a sum of a set of functions of the factors that influence spindles, each multiplied by a model parameter to be estimated:

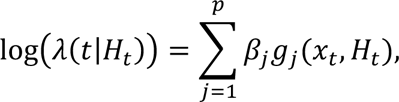

where *p* is number of model parameters, *β*_j_ is the *j*^th^ model parameter, and *g*_j_(*x*_t_, *H*_t_) are a set of functions of the variables, *x*_t_, that influence the intensity of events and *H*_t_, that determine how past history influences the intensity of events. The specific form of each model used to analyze spindle data in this paper is provided in table 1.

**Table 1:**
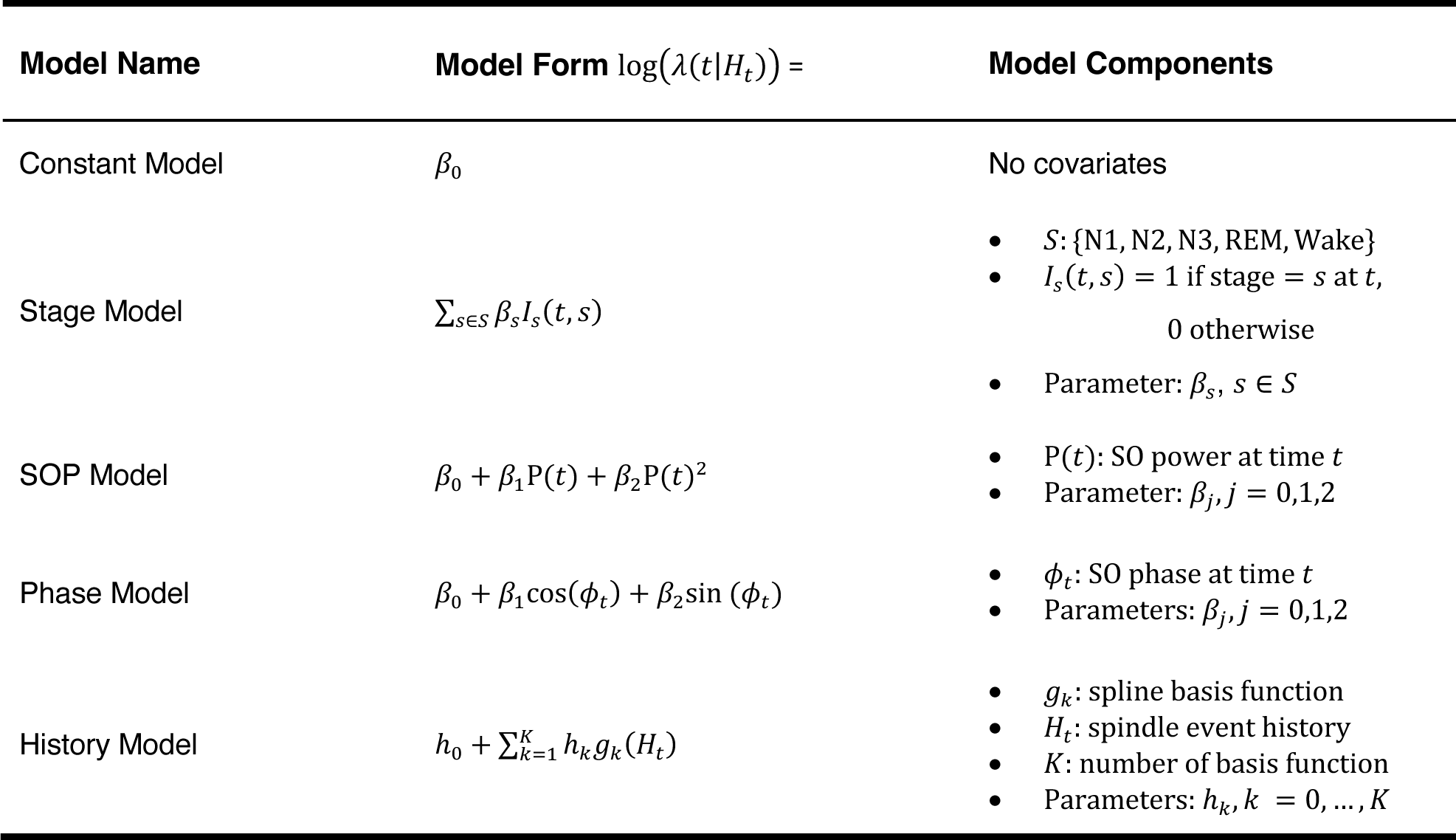
Spindle dynamics model forms.

| Model Name | Model Form $\log(\lambda(t\vert H_t)) =$ | Model Components |
| --- | --- | --- |
| Constant Model | $\beta_0$ | No covariates |
| Stage Model | $\sum_{s \in S} \beta_s I_s(t, s)$ | $S: \{N1, N2, N3, REM, Wake\}$<br>$I_s(t, s) = 1$ if stage = $s$ at $t$,<br>0 otherwise<br>Parameter: $\beta_s, s \in S$ |
| SOP Model | $\beta_0 + \beta_1 P(t) + \beta_2 P(t)^2$ | $P(t)$: SO power at time $t$<br>Parameter: $\beta_j, j = 0, 1, 2$ |
| Phase Model | $\beta_0 + \beta_1 \cos(\phi_t) + \beta_2 \sin(\phi_t)$ | $\phi_t$: SO phase at time $t$<br>Parameters: $\beta_j, j = 0, 1, 2$ |
| History Model | $h_0 + \sum_{k=1}^K h_k g_k(H_t)$ | $g_k$: spline basis function<br>$H_t$: spindle event history<br>$K$: number of basis function<br>Parameters: $h_k, k = 0, \dots, K$ |

This model framework allows us to test which of multiple factors influences the instantaneous spindle rate, estimate the functional form of these influences, and determine how these influences interact. In table 1, we list a number of possible single predictors including sleep stage, spindle past history, SO phase, and SO power. We then build up a set of models by sequentially combining multiple predictors and including their interactions to create larger and more detailed models. The full list of model forms is provided in supplementary materials.

### 4.4. Model inference methods

Point process GLMs have a number of properties that make them useful for the analysis of spindle event data. They can estimate the simultaneous influences of multiple factors, including non-linear influences, with interpretable parameters^44,95^. These models produce convex likelihoods enabling computationally efficient algorithms (e.g., iteratively reweighted least squares (IRLS)) to compute the maximum likelihood estimators for GLM parameters^44,51,96,97^. These algorithms are supported by robust software packages in popular statistical platforms and coding languages such as Python, MATLAB, and R. The fitted parameters from maximum likelihood estimation are asymptotically normal, unbiased, and have minimum mean-squared error. Additionally, the Fisher information for the parameters is computed from the IRLS algorithm, which determines the standard errors of the model parameters, and formal statistical tests can be conducted to determine whether any collection of variables significantly influence the event intensity^97^.

Point process GLMs also provide well-established tools for assessing model goodness-of-fit. The model deviance explained is an analogue of variance explained in linear regression, which can be used to compare models^51^. Another goodness-of-fit tool is based on the time-rescaling theorem, which transforms the observed inter-spindle intervals into the wait times for a homogenous Poisson process if the model is correct^65^. After rescaling, Kolmogorov-Smirnov (KS) plots are used to compare the distribution of inter-spindle times to those predicted by the model. A well-fitted model produces a KS plot closely following a 45-degree line, and everywhere contained within its significance bounds, while deviations outside the bounds suggest model misfit. A KS test compares the largest deviation between the empirical and model distributions to known critical values for the hypothesis that the model is correct.

### 4.5. Artifact detection

Artifact detection is implemented in the time domain by an iterative procedure described in Stokes et al^2^. Briefly, to detect high-frequency artifacts, the raw data is filtered into a high-frequency (35 Hz to Nyquist) component, and then any data outside of the threshold (mean ± 3.5 × standard deviation) is identified as an artifact and removed; To detect noise with broadband energy, the raw data is filtered from 2 Hz to Nyquist, then the same exclusion criteria are applied to mark artifacts. This iterative approach computes a new threshold each time and removes artifacts until no data exceeds the threshold.

### 4.6. Event detection

The automated spindle detection method is based on our previous work^1^, which identifies transient oscillation activity using time-frequency peaks in the sigma range of the EEG spectrogram, which are termed TFσ peaks. Traditionally-scored spindles have been shown to be but a subset of the underlying neurophysiological activity within the spindle range of the EEG^1,2^, due to the origins of spindle identification based on time domain identification by eye based on standards from the 1930s^4^. This approach has been shown to provide a less biased representation of the spindle activity with increased night-to-night stability and greater statistical power than traditional spindle identification approaches while retaining the equivalent waveform morphology^1^. To investigate fast spindle dynamics, we detect TFσ peaks in the frequency range from 12 to 16 Hz, commensurate with the “fast” spindle activity that is primarily studied. The detected peak time-frequency centroids were used as the timing for point process events. Event times were then discretized into 0.1-second intervals, and a binary spindle event train was computed, indicating whether a spindle event (0 or 1) occurred in each time bin. For simplicity, we refer to the events as spindles, but it is vital to note we are using a more robust and principled event basis. Implementation details are provided in the supplementary materials.

### 4.7. Slow oscillation phase and power

To estimate the slow oscillation phase (SO-phase), the EEG signal was first band passed from 0.4 to 1.5 Hz using a zero-phase filter, and the Hilbert transform of the filtered signal was applied to create the analytic signal. The instantaneous phase was estimated by computing the angle of the analytic signal. TFσ peak phase was computed using linear interpolation of the unwrapped phase at the TFσ peak times (defined by the spectral peak centroids), and all the phase values were re-wrapped to −*π* to *π*, where phase 0 means the peak of the slow oscillation, −*π* to 0 and 0 to *π* correspond to rising and falling phase, respectively.

To estimate the slow oscillation power (SO-power), a multitaper spectrogram^98,99^ of the EEG signal after artifact detection was computed with the following parameters: 3 tapers and time-half bandwidth of 2, 4-second window length with 1-second step size. The raw SO-power was then computed by integrating the power in the slow oscillation frequency range (0.4 – 1.5 Hz) and converting it to decibels (dB). To standardize across participants, the SO-power was normalized by subtracting the 2^nd^ percentile of the raw SO-power in non-wake stages (REM + NREM), which tends to coincide with light or N1 sleep. This acts as an objectively defined point of alignment for comparison across individuals and populations.

### 4.8. History modulation curve summary statistics

Given the variability in history dependence structure observed across the population, we explored a set of summary statistics that allow us to capture and interpret the temporal patterns. Based on the history modulation curve, as well as its 95% upper confidence bound (UCB) and lower confidence bound (LCB) at each time lag, we define the following summary statistics:

1. Refractory period – The first lag at which the UCB is greater than or equal to 1.
2. Excitatory period – The length of the period from when the LCB is first greater than or equal to 1 to when the LCB returns to be less than or equal to 1.
3. Peak height – The maximum value of the history modulation.
4. Peak time – The lag corresponding to peak height.

### 4.9. Cardinal spline functions

History dependence was fit with a GLM as a function of history using cardinal spline basis functions, which have been successfully employed to capture temporal patterns in neural spiking activity^100,101^ and sleep respiratory data^46^. We set a tension parameter of 0.5, with end points at 0 and 15 seconds, six knots were evenly placed in every 1.5 seconds up to 9 seconds, with another knot at 12 seconds. Two additional knots placed at −10 and 25 seconds were used to determine the derivatives of the spline function at the end points. For a detailed overview of incorporating spline models into a GLM framework, see Sarmashghi et al^100^.

### 4.10. Preferred phase and confidence interval

All the models were fit using the GLM package in MATLAB. The maximum likelihood estimator for the preferred phase was computed from the GLM parameter fits. For example, after fitting in the *Phase Model* (Figure 7d), the estimated preferred phase is the four-quadrant inverse tangent^102^ of 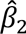 and 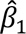. Confidence intervals for the preferred phase were computed as follows. Since the fitted model parameters asymptotically follow a multivariate normal distribution (MVN)^44,51^, 10000 random samples of 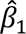 and 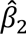 were drawn from 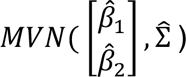, where 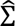 is the estimated covariance matrix provided by the GLM fit. The estimated preferred phase was computed for each sample and the 2.5^th^ and 97.5^th^ quantiles were computed. The preferred phase and confidence interval for other models that included the SO phase component were derived similarly. In Figure 6c, the mean parameters from the *Phase-SOP Model* in each age group were used to visualize the mean phase shift curve.

### 4.11. Deviance explained

The mean fractional deviance explained across MESA is reported in Result section 2.9, which represents the fraction of the total deviance explained by the fully model that is explained by each individual factor. We first computed the total deviance reduction (Dev_null_ – Dev_full_) by comparing the null *Constant Model* to the full *Stage-Phase-History Model,* then computed the deviance reduction for the stage component (Dev_null_ – Dev_stage_) by comparing the null *Constant Model* to the *Stage Model*. The fractional deviance explained is defined as 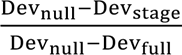. Deviance explained by phase, SOP and history are computed similarly.

### 4.12. Synergy index

We adapt a commonly used measure of information independence for neural coding models^66–69^, which is defined as the joint information (*I_AB_*) provided by two factors, A and B, minus the sum of the information provided by A and B alone (*I_A_* + *I_B_*). A normalized synergy measure^66^ is 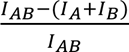. It ranges from −1 to 1, where −1 indicates A and B provide totally overlapping (redundant) information, 0 indicates A and B are independent, and 1 indicates that while A or B alone provides no information, they jointly convey complete (synergistic) information.

To test whether phase and history interact to influence spindle dynamics (Figure 8b), we computed the information as the deviance explained by each model, so that the synergy index was 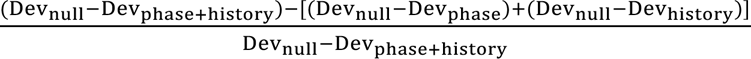, where Dev_phase_ and Dev_history_ are the deviance of the models that include the phase, and history components alone, respectively, and Dev_phase+history_ is the deviance from the model that includes both of these factors.

### 4.13. Statistical tests

Maximum likelihood ratio tests were performed to determine whether spindle history, phase, and stage-dependent phase coupling contributed significantly to spindle intensity. Kolmogorov-Smirnov (KS) tests were used to evaluate the goodness-of-fit of the models (Figure 7e, KS pass rate in Result section 2.9). In Figure 4b & Figure 6c, permutation tests with global bounds were performed to compare history curves across different groups^103^. t-tests (uncorrected for multiple comparison) were conducted in Figure 3b & Figure 4a to compare the mean rates (∗, ∗∗, ∗∗∗ denote the p-value <.05, <.01, <.001 separately). In Figure 6a & b, a Watson-Williams test was performed to test the significance of the preferred phase shift across sleep depth. All statistical analyses were performed in MATLAB_R2022b. Significance levels of 0.05 were used, if not otherwise specified.

## CODE AVAILABILITY

Code for the analysis in this paper is available at http://sleepEEG.org.

## DATA AVAILABILITY

The Multi-Ethnic Study of Atherosclerosis (MESA)^49,94^ dataset was obtained from the National Sleep Research Resource. Two-night data from Wamsley et al^19^ was obtained by permission from Dr. Dara Manoach.

## AUTHOR CONTRIBUTIONS

M.J.P., U.T.E., and S.C. designed the study. S.C. and M.H. preprocessed the data. S.C., M.J.P., and U.T.E. performed the statistical modeling. S.C. drafted the initial manuscript. M.J.P. and U.T.E. supervised the work. All authors reviewed, revised, and approved the final version of the manuscript.

## ACKNOWLEDGEMENTS

The authors would like to express their deep gratitude towards Dr. Dara Manoach for access to the two-night data from Wamsley et al. The Multi-Ethnic Study of Atherosclerosis (MESA) Sleep Ancillary study was funded by NIH-NHLBI Association of Sleep Disorders with Cardiovascular Health Across Ethnic Groups (RO1 HL098433). MESA is supported by NHLBI funded contracts HHSN268201500003I, N01-HC-95159, N01-HC-95160, N01-HC-95161, N01-HC-95162, N01-HC-95163, N01-HC-95164, N01-HC-95165, N01-HC-95166, N01-HC-95167, N01-HC-95168 and N01-HC-95169 from the National Heart, Lung, and Blood Institute, and by cooperative agreements UL1-TR-000040, UL1-TR-001079, and UL1-TR-001420 funded by NCATS. The National Sleep Research Resource was supported by the National Heart, Lung, and Blood Institute (R24 HL114473, 75N92019R002).

## SUPPLEMENTARY MATERIAL

Supplementary material is provided in a separate document and will be available online.

