## Supplementary Material for "Individualized temporal patterns dominate cortical upstate and sleep depth in driving human sleep spindle timing"

**S1 Spindle detection algorithm**

This automated spindle detection method is based on our previous work from Dimitrov et al.^1^, which extracts and identifies time-frequency peaks in the sigma range of the EEG spectrogram (TFσ peaks). The goal of this approach is to create a method of spindle detection that is not as prone to the amplitude selection biases and rarity assumptions present in traditional spindle detection. In this section, we outline an enhanced version of the approach that demonstrates greater robustness and accuracy.

**Event extraction from the time-frequency domain**

The details of this event extraction are described explicitly in Dimitrov et al. Everything is preserved except for the clustering algorithm, which we will demonstrate in the next section. However, here is an overview of the algorithm.

Our method focuses on analyzing transient oscillatory activity in the time-frequency domain, which appears as salient peaks in the multitaper spectrogram. The parameter settings for the multitaper spectrogram are as follows: a 1-second window with a 0.05-second step size, 3 tapers, and a time-half bandwidth of 2. This algorithm extracts these peaks by searching for regions with distinct peak-like structures in both time and frequency dimensions. This is done in two steps, using the concept of peak prominence, which measures the height of a peak relative to its local baseline.

In the first step, called the "frequency step," we detect peaks in the EEG power spectrum at each time and estimate the prominence of the largest peak in the spindle frequency range. In the second step, the "time step," we identify temporal peaks in the time trace of prominence values obtained from the "frequency step." Each temporal peak found in the "time step" therefore corresponds to a well-defined local maximum on the spectrogram. To describe the shape of these time-frequency peaks, we calculate properties such as prominence, duration, central frequency, and bandwidth (quantified in Dimitrov supplementary materials). Before clustering to separate noise peaks, we exclude peaks with durations of less than 0.3 seconds to match established criteria. Additionally, we exclude peaks with frequency bandwidths less than half of the spectral resolution of the spectrograms, as they are not accurately measurable.

**Updated clustering algorithm**

The original algorithm used a two-class k-means clustering on the prominence values of all time-frequency local maxima in the 9–17 Hz range. However, this approach is effective only when there is a clear separation between actual signals (i.e., putative TFσ peaks) and noise events. The original algorithm is robust for healthy young adults, where spindles and TFσ peaks are distinct, and signal events have noticeable prominence values that can be separated from the distribution of noise events. Yet, in other groups, particularly older adults where TFσ peaks are less pronounced, the previous method does not perform well because the prominence distribution is unimodal, making the two-class k-means clustering ineffective in automatically identifying TFσ peaks.

To address this challenge and improve the algorithm's general applicability, we analyzed the distributions of four event properties extracted from spectrograms: prominence, central frequency, duration, and bandwidth. Despite variations in the strength of spindles and TFσ peaks, the duration and bandwidth properties consistently exhibit separable peaks at higher values. These properties consistently show triplet peak structures, likely corresponding to the split of noise local maxima during spectral estimation.

Instead of relying on a single two-class k-means clustering on log prominence, which could inaccurately split the large noise peak in half, we now apply two separate three-class clusterings on duration and bandwidth. The cluster centers are seeded at 0 sec, 0.3 sec, and 1 sec for duration, and at 0.5 Hz, 2 Hz, and 3.5 Hz for bandwidth. The final separation of signal and noise is determined by combining the cluster memberships from these two properties. Specifically, a time-frequency local maximum is identified as a TFσ peak if its duration and bandwidth both belong to the largest clusters on the two dimensions. This enhanced method for detecting TFσ peaks offers better distinction between signal and noise and is currently the default setting in this algorithm.

The code for the spindle detection algorithm is available as a component of the DYNAM-O toolbox, available at <http://sleepEEG.org>.

**S2 Full list of model forms with interaction terms**

We list all model forms that are used for addressing specific questions of spindle temporal dynamics.

| **Model Name** | **Model Form** $\log\left( \lambda\left( t \vert H_{t} \right) \right)$ = | **Model Components** |
| --- | --- | --- |
| Constant Model | $\beta_{0}$ | No covariates |
| Stage Model | $\sum_{s\in S} \beta_{s}I_{s}\left( t,s \right)$ | - $S:$ {N1, N2, N3, REM, Wake} - $I_{s}\left( t,s \right)=1 if stage=s\mathrm{at}t,$   $0 otherwise$   - Parameters: $\beta_{s}$, $s\in S$ |
| SOP Model | $\beta_{0}+\beta_{1}P(t)+\beta_{2}{P(t)}^{2}$ | - $P(t$): SO power at time $t$ - Parameter: $\beta_{j}, j=0,1,2$ |
| Phase Model | $\beta_{0}+\beta_{1}\cos\left( \phi_{t} \right)+\beta_{2}sin(\phi_{t})$ | - $\phi_{t}$: SO phase at time $t$ - Parameters: $\beta_{j}, j=0,1,2$ |
| History Model | $h_{0}+\sum_{k=1}^{K} h_{k}g_{k}\left( H_{t} \right)$ | - $g_{k}:$ spline basis function - $H_{t}:$ spindle event history - $K:$ number of basis function - Parameters: $h_{k}, k =0,\ldots,K$ |
| Stage-History Model | $\sum_{s\in S} \beta_{s}I_{s}\left( t,s \right)+\sum_{k=1}^{K} h_{k}g_{k}\left( H_{t} \right)$ | Parameters: $\beta_{s}$, $s\in S; h_{k}, k =1,\ldots,K$ |
| Phase-Stage Model | $\beta_{0}+\beta_{1}\cos\left( \phi_{t} \right)+\beta_{2}\sin\left( \phi_{t} \right)+\sum_{s\in S} \beta_{s}I_{s}\left( t,s \right)+\sum_{s\in S} \alpha_{s}I_{s}\left( t,s \right)\cos\left( \phi_{t} \right)+\sum_{s\in S} \gamma_{s}I_{s}\left( t,s \right)\sin\left( \phi_{t} \right)$ | Parameters: $\beta_{j}, j=0,1,2; \beta_{s}$, $s\in S; \alpha_{s}$, $s\in S ; \gamma_{s},s\in S$ |
| Phase-SOP Model | $\beta_{0}+\beta_{1}P\left( t \right)+\beta_{2}{P\left( t \right)}^{2}+\beta_{3}\cos\left( \phi_{t} \right)+\beta_{4}\sin\left( \phi_{t} \right)+\beta_{5}P\left( t \right)\cos\left( \phi_{t} \right)+\beta_{6}P\left( t \right)\sin\left( \phi_{t} \right)+\beta_{7}{P\left( t \right)}^{2}\cos{(\phi}_{t})+\beta_{8}{P\left( t \right)}^{2}\sin{(\phi}_{t})$ | Parameters: $\beta_{j}, j=0,1,\ldots,8$ |
| Phase-History Model | $\beta_{0}+\beta_{1}\cos\left( \phi_{t} \right)+\beta_{2}\sin\left( \phi_{t} \right)+\sum_{k=1}^{K} h_{k}g_{k}\left( H_{t} \right)$ | Parameters: $\beta_{j}, j=0,1,2; h_{k}, k =1,\ldots,K$ |
| Stage-Phase-History Model | $\beta_{0}+\beta_{1}\cos\left( \phi_{t} \right)+\beta_{2}\sin\left( \phi_{t} \right)+\sum_{s\in S} \beta_{s}I_{s}\left( t,s \right)+\sum_{k=1}^{K} h_{k}g_{k}\left( H_{t} \right)$ | Parameters: $\beta_{j}, j=0,1,2; \beta_{s}$, $s\in S; h_{k}, k =1,\ldots,K$ |
| SOP-Phase-History Model | $\beta_{0}+\beta_{1}P\left( t \right)+\beta_{2}{P\left( t \right)}^{2}+\beta_{3}\cos\left( \phi_{t} \right)+\beta_{4}\sin\left( \phi_{t} \right)+\sum_{k=1}^{K} h_{k}g_{k}\left( H_{t} \right)$ | Parameters: $\beta_{j}, j=0,1,\ldots,4;$ $h_{k}, k =1,\ldots,K$ |
| Stage-Phase-History Interaction Model | $\beta_{0}+\beta_{1}\cos\left( \phi_{t} \right)+\beta_{2}\sin\left( \phi_{t} \right)+\sum_{s\in S} \beta_{s}I_{s}\left( t,s \right)+\sum_{s\in S} \alpha_{s}I_{s}\left( t,s \right)\cos\left( \phi_{t} \right)+\sum_{s\in S} \gamma_{s}I_{s}\left( t,s \right)\sin\left( \phi_{t} \right)+\sum_{k=1}^{K} h_{k}g_{k}\left( H_{t} \right)$ | Parameters: $\beta_{j}, j=0,1,2; \beta_{s}$, $s\in S; \alpha_{s}$, $s\in S ; \gamma_{s},s\in S$; $h_{k}, k =1,\ldots,K$ |
| SOP-Phase-History Interaction Model | $\beta_{0}+\beta_{1}P\left( t \right)+\beta_{2}{P\left( t \right)}^{2}+\beta_{3}\cos\left( \phi_{t} \right)+\beta_{4}\sin\left( \phi_{t} \right)+\beta_{5}P\left( t \right)\cos\left( \phi_{t} \right)+\beta_{6}P\left( t \right)\sin\left( \phi_{t} \right)+\beta_{7}{P\left( t \right)}^{2}\cos{(\phi}_{t})+\beta_{8}{P\left( t \right)}^{2}\sin{(\phi}_{t})+\sum_{k=1}^{K} h_{k}g_{k}\left( H_{t} \right)$ | Parameters: $\beta_{j}, j=0,1,\ldots,8$; $h_{k}, k =1,\ldots,K$ |

**S3 Supplementary Figure List**

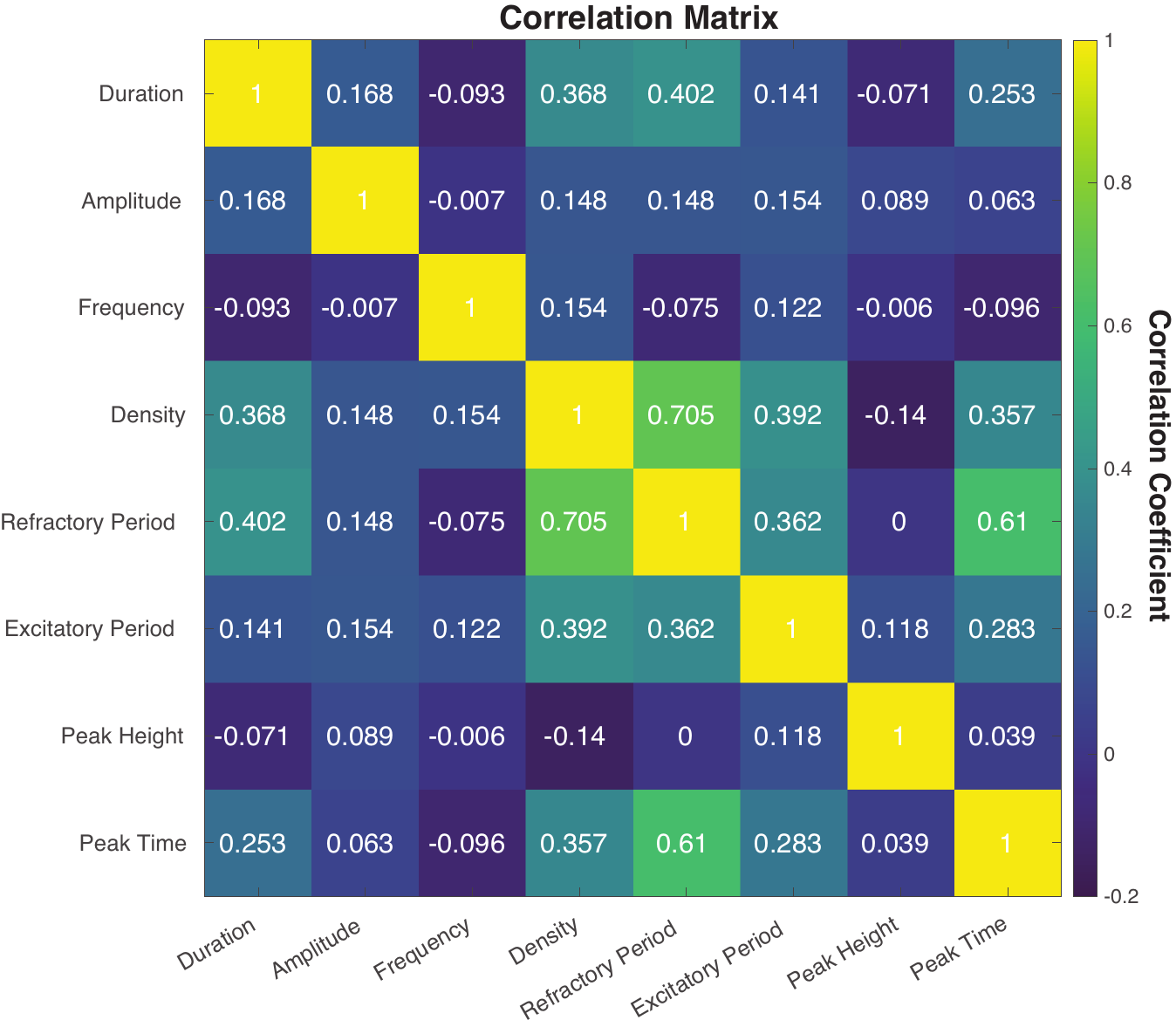

**Supp Figure 1: Mean correlation matrix for spindle history features and spindle morphologies across MESA population.**

**Supp Figure 2: Mean correlation matrix for different modeling components across MESA population.**

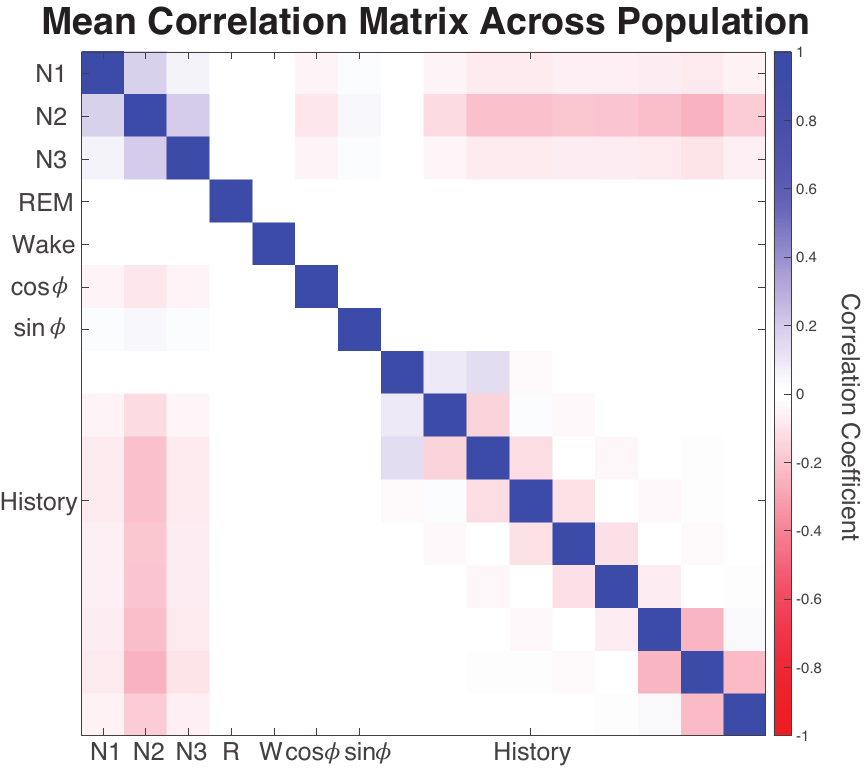
